# Multimodal and multisensory coding in the *Drosophila* larval peripheral gustatory center

**DOI:** 10.1101/2020.05.21.109959

**Authors:** G. Larisa Maier, Marjan Biočanin, Johannes Bues, Felix Meyenhofer, Clarisse Brunet Avalos, Jae Young Kwon, Bart Deplancke, Simon G. Sprecher

## Abstract

The ability to evaluate food palatability is innate in all animals, ensuring their survival. The external taste organ in *Drosophila* larvae is composed of only few sensory neurons but enables discrimination between a wide range of chemicals and displays high complexity in receptor gene expression and physiological response profile. It remains largely unknown how the discrepancy between a small neuronal number and the perception of a large sensory space is genetically and physiologically resolved. We tackled dissection of taste sensory coding at organ level with cellular resolution in the fruit fly larva by combining whole-organ calcium imaging and single-cell transcriptomics to map physiological properties and molecular features of individual neurons. About one third of gustatory sense neurons responded to multiple tastants, showing a rather large degree of multimodality within the taste organ. Further supporting the notion of signal integration at the periphery, we observed neuronal deactivation events within simultaneous neighboring responses, suggesting inter-cellular communication through electrical coupling and thus providing an additional level in how neurons may encode taste sensing. Interestingly, we identified neurons responding to both mechanical and taste stimulation, indicating potential multisensory integration. On a molecular level, chemosensory cells show heterogeneity in neuromodulator expression. In addition to a broad cholinergic profile, markers on dopaminergic, glutamatergic or neuropeptidergic pathways are present either in distinct cell populations or are seemingly co-expressed. Our data further extend the sensory capacity of the larval taste system pointing towards an unanticipated degree of multimodal and multisensory coding principles.

## Introduction

Taste sensing provides the innate ability in all animals to avoid toxic or deleterious food ingestion and to choose a nutritious diet. The fruit fly larva is able to discriminate between a large range of different gustatory cues (1), being – similarly to humans – repelled by bitter tastants (2-4) as possible signals of toxic food, and attracted by sweet and certain amino acids (5-7), indicators of a nutritious diet. Distinct from mammals where hundreds of taste cells are assembled in taste buds on the tongue, the larva disposes of very few gustatory sense neurons (GSNs) at the periphery (8, 9), raising the question of how does a small neuronal population encode a large palette of cues.

Distinct models of sensory coding have emerged for taste perception. According to the labeled-line model, each cell within a mammalian taste bud is tuned to one of the 5 basic tastes – sweet, umami, bitter, sour and salt (10-13). This model relies to a large extent on the existence of a best stimulus concentration (14) but varying the concentration of a given tastant alters the number of responding cells and afferent sensory neurons, often increasing the proportion of multituned neurons (15). Moreover, taste cells within a bud have been shown to communicate with each other through purinergic signaling (16), entailing coexistence of narrowly and broadly tuned taste sensing cells (17) and signal integration at the periphery, distinct from a strictly segregated taste sensing model.

In *Drosophila* taste sensing has been described in accordance with a labeled-line model, as sweet, bitter and water perception localize in separate populations of peripheral GSNs (18-22). However, additional tastes such as salt, fatty acid, carbonation, polyamines or amino acids partially overlap onto sweet or bitter sensing GSNs by means of specific added-on receptor expression (23-30), suggesting a model of taste sensing with both narrowly and more broadly tuned cells, similar to mammals.

The peripheral gustatory system in the *Drosophila* larva presents many similarities with the adult fly, such as the sensillar organization of dendrites, presence of both internal and external chemosensory organs (31), as well as cue detection through the same chemoreceptor gene families – GRs (Gustatory Receptors), IR (Ionotropic Receptors), PPKs (pickpocket) or TRPs (Transient Receptor Potential) (29, 32-38). On the other hand, larval taste processing presents developmental stage particularities, considering that the number of GSNs is much lower than in adult flies and many receptors are larva-specific (39, 40).

The main chemosensory organs in *Drosophila* larva are located at the tip of the head, represented by ganglia of bipolar sensory neurons (Fig.1A) (41). External organs include the terminal organ ganglion (TOG) – with main taste, but also mechano- and thermo-sensory function, the dorsal organ ganglion (DOG) – with primary olfactory role, and the smaller, generally uncharacterized ventral organ ganglion (VOG) – with associated taste function (8). Sensory neurons in these ganglia extend dendrites to the periphery in direct contact with the environment and axonal projections into defined regions of the brain. Conversely, internal chemosensory organs called ventral, dorsal and respectively posterior pharyngeal sensories (VPS, DPS, PPS) are located on the pharyngeal tube where they project their dendrites (42), therefore functionally associated with taste sensing during food ingestion.

Thanks to the 1 to 1 receptor to neuron expression in the DOG, functional dissection of the 21 olfactory sensory neurons was achieved using corresponding Gal4 lines (43-47). In contrast, the neighboring TOG, main external larval taste center, comprises more than 30 GSNs out of which only 7 neuronal identities (C1-C7) have been mapped and can be traced using individual Gal4 lines (Fig.1B) (39, 48, 49). Characterization of single larval GSNs has revealed multimodality and even taste integration of opposed valence within one GSN (48), but the general taste sensing logic in this animal remains largely unexplored.

We here delve into sensory tuning diversity of TOG neurons by using whole-organ in-vivo calcium imaging recordings on larvae expressing genetically encoded calcium sensors. To gain insight into molecular components of TOG and DOG neurons we employed DisCo, a novel approach that enables single-cell RNA sequencing of low input samples (50). Our findings provide insights into fundamental aspects of taste sensory coding. First, we show that roughly one third of responding GSNs are activated by more than one taste modality, while conversely, single tastants generally elicit responses in several neurons. Interestingly, we detected neuronal deactivation responses, distinct from canonical calcium imaging signals, which occurred within concomitant firing in adjacent neurons, as described for electrical coupling in the adult fly olfactory system (51, 52). Using DisCo and immunostainings, we identified distinct and common molecular markers in TOG/DOG neurons. While it was known that larval chemosensory neurons are cholinergic (53), we found that they are also glutamatergic and dopaminergic, and, similar to the central nervous system (CNS) (54), neurons may also co-express neurotransmitter markers. Lastly, we detected expression of mechanosensory receptor genes and, intriguingly, GCAMP recordings showed activation to both mechanical and chemical stimulation in some neurons, implying multisensory integration at single cell level. Taken together, our observations of the larval external gustatory organ bring evidence of multimodality, possibly inter-cellular communication through electric coupling, heterogeneous expression of neurotransmitter markers and mechanosensory features in GSNs.

## Results

### Whole organ larval taste recordings with single cell resolution

Due to the sparse availability of Gal4 lines for individual GSN labeling in the larva, assessing an overview of peripheral taste coding by tackling single neurons entails limitations. We therefore undertook characterization of TOG neurons by means of a whole-organ approach.

First, we performed immunostainings on individual GSNs expressing *myr*:*GFP* (Fig.1C & D). Structural flexibility of this tissue entails variability across samples but we noted the 7 previously identified GSNs maintained a relatively stereotypical localization between stainings (n≥3) (Fig1C). This allowed assembling 3D locations of the 7 GSNs, serving as approximation map of their respective position within the organ (Fig.1D).

Further, we performed whole-organ calcium imaging recordings by expressing cytoplasmic GCaMP6m (55) and nuclear RFP (56) reporters in all neurons driven by *nSyb*-Gal4 (Fig.1E). The use of a nuclear reporter as cellular landmark was critical for whole organ recordings, as solely a cytoplasmic fluorophore would not sufficient for efficient segmentation due to low baseline fluorescence intensity in some cells and to ambiguous delimitation between neighboring somata. As previously described (57), we made use of a customized microfluidic chamber for chemical stimulation while recording physiological responses of all GSNs in a semi-intact larval preparation, and we subsequently developed a data processing pipeline for whole organ recordings (Suppl.Fig.1).

### One third of responding GSNs show taste multimodality

We first tested two distinct series of chemicals each containing 5 substances belonging to the 5 canonical taste categories: sweet, bitter, sour, amino acids and salt. Substances and concentrations were chosen according to former studies (48, 58) (Table1, Fig.2A, see Methods). Out of all responding neurons across 15 total recorded organs, 66% showed activation to only one tastant per animal and most unimodal responses were recorded to sucrose and high salt (Fig.2A, uni-taste/cell). In line with our previous findings that some GSNs are multimodal (48), the remaining 34% of responding neurons were activated by more than one substance (Fig.2A, Suppl.File1), showing that several larval GSNs have the capacity to respond to different taste categories. The top 5 taste response combinations mapping to one neuron (Fig.2A, multi-taste/cell) were: sucrose and valine/arginine (sweet + amino acid, 17%), sucrose and citric acid (sweet + sour, 17%), citric acid and high salt (sour + high salt, 14%), sucrose and high salt (sweet + high salt, 14%) and finally sucrose and denatonium/quinine (sweet + bitter, 14%). Interestingly, sucrose was the tastant corresponding to most uni-taste/cell responses but also to most frequent taste combinations of multi-taste/cell.

Next, we expanded the number of gustatory cues used for GSN stimulation by using groups of tastants for sweet, bitter, amino acid and salt taste categories (Fig.2B, Suppl.Table 1). To begin with, we tested 2 groups of sugars, chosen as monosaccharides (fructose, glucose, arabinose, mannose, galactose), and respectively disaccharides (sucrose, trehalose, maltose, lactose, cellobiose). We calculated a rounded 74% of sugar-responsive neurons to be activated by either monosaccharides (41%) or disaccharides (33%), and a rounded 25% by both groups (Fig.2B, upper left panel).

For salt taste stimulation we mixed NaCl and KCl for a final concentration of 50mM for low salt group and respectively 1M for high salt group. We observed a low percentage of cell activation overlap among salt-responding cells, 20% being activated by both groups while the rest responded either to high salt (56%) or to low salt (24%).

The 20 proteinogenic amino acids were compiled in 4 groups (A, B, C and D) as previously described by Park et al. (59) (Suppl.Table 1). 38% amino acid-responding neurons showed activation to 2 or more amino acid groups. Similar percentages were observed for animals stimulated with all 8 groups (2 sugar, 2 salt and 4 amino acid units), with up to 40% neurons responding to at least 2 different taste modality groups (Fig.2B upper right panel).

Bitter tastants were randomized in 2 groups of 4 substances each (Suppl.Table 1): DSoTC group (denatonium benzoate, sucrose octaacetate, theophylline, coumarin), and respectively QLSC group (quinine, lobeline, strychnine, caffeine). Among cells responsive to bitter stimulation we observed a bigger fraction of cells activated by DSoTC group (55%) than by QLSC (30%). In animals tested for sweet and bitter taste response, the percentage of cells activated by both sugar and bitter groups was of 30% (Fig.2B, lower panels), in agreement with cell integration of tastants with opposite valence.

### Taste stimulation can elicit either activation or deactivation signals in GSNs

While canonical responses of GSNs to taste cues entail a rise of GCaMP fluorescence, we also detected neuronal responses characterized by a fluorescence intensity decrease. For simplicity, and not as conclusive categorization, we named these observed intensity fluorescence decrease signals: *deactivation responses*, as they are seemingly opposite to canonical neuronal activation measured through fluorescence increase.

A notable example of deactivation was recorded in response to citric acid in 2 neurons localized centrally in the organ in close proximity with each other. We termed these neurons CDL1 (central-dorsal-lateral neuron 1) and CDL2, respectively. Intriguingly, CDL1 and CDL2 responded to citric acid in a pattern of synchronous activation-deactivation (Fig.3A).

To examine into the possibility that deactivation responses were caused by top-down inhibition by central interneurons, we performed recordings in animals with severed connections between chemosensory neurons and CNS (Fig.3B). Detaching the brain and the respective sensory nerve connections did not abolish the activation-deactivation responses induced by citric acid stimulation. Likely, this pattern corresponds to ephaptic transmission where the electrical field of a responding cell generates hyperpolarization in the partner cell (60), also reported in the adult olfactory system (51).

### Mapping neuronal responses across larvae shows a high level of taste multimodality

Next, we proceeded to match neurons based on their spatial position within the TOG organ and responses recorded to individual taste series presented in figure 2A. We identified and named 21 GSNs (Fig.4, y axis and 3D image). All considered responses exceeded 20% DF/F0 and were manually validated. Not all neurons showed the same response consistency, as reflected by the stereotypy of response, meaning the percentage of times a neuron was identified to respond (Fig.4A freq, grey tones with darker color denoting higher frequency). Heatmap for activation and deactivation responses for each neuron were compiled separately and for each cell and each tastant, we plotted the stereotypy of response next to the average intensity of response (Fig.4A, x axis). Average intensities were considered only if response frequencies were at least 20% for the given tastant.

**Figure 1:**
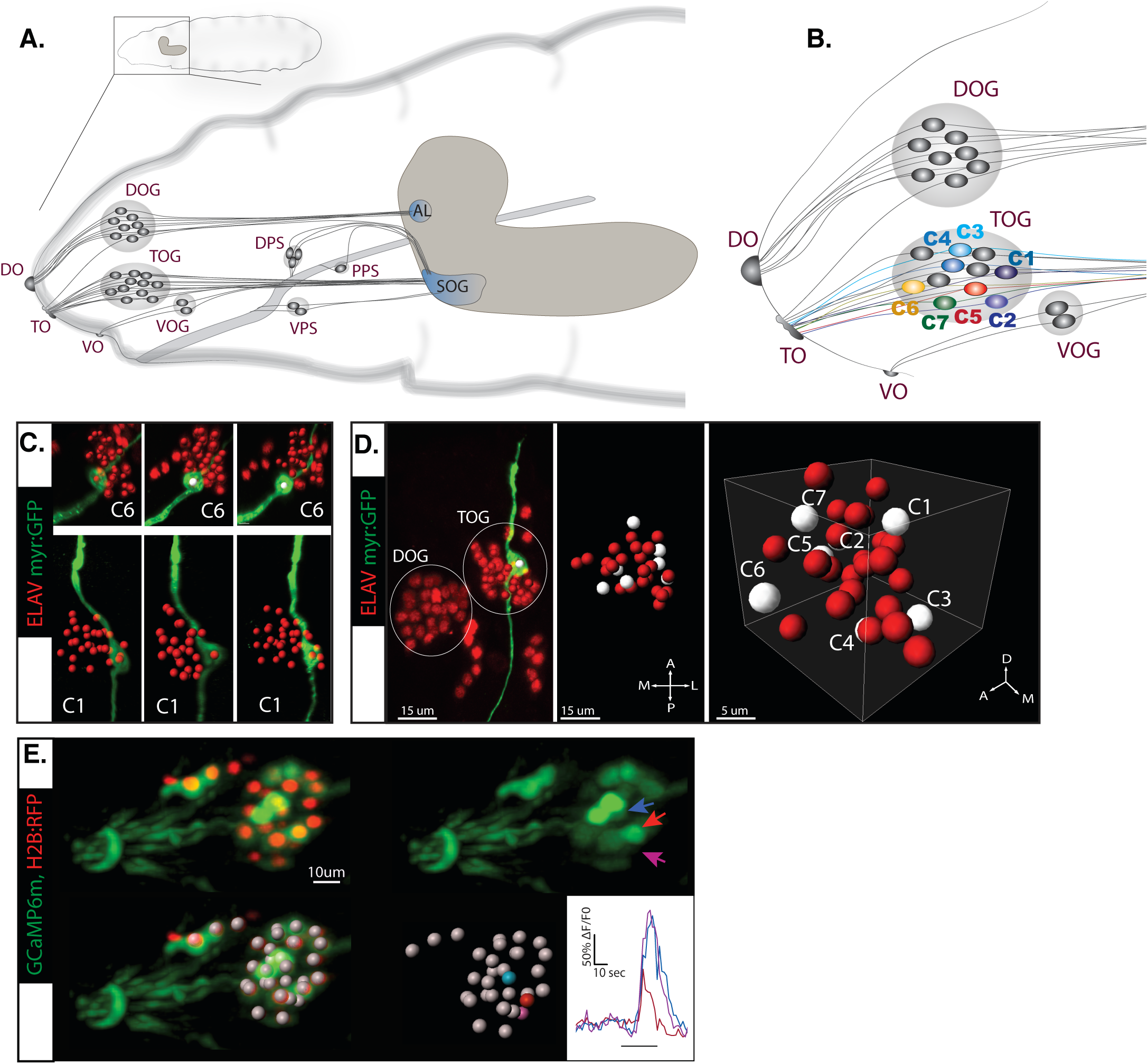
Peripheral taste neurons of the fruit fly larva – visualization and calcium imaging. A. The chemosensory system of the larva. External sense organs extend dendrites to the periphery and have main olfactory function – the DOG (dorsal organ ganglion), or gustatory function – the TOG (terminal organ ganglion) and the VOG (ventral organ ganglion). Dendrites of the internal dorsal, ventral and posterior pharyngeal sense organs (DPS, VPS and PPS) innervate the pharynx, thus involved in taste sensing during food ingestion. All chemosensory neurons project axons to the brain in the subesophageal ganglion (SOG) – first central taste integration relay – or to the antennal lobe (AL) for central olfactory processing. Adapted from (1). B. External chemosensory organs. 7 GSNs previously identified and named C1-C7 are represented here by the color code they were first described with (39, 48). Used Gal4 lines: *Gr22e*-Gal4 (C1), *Gr94a*-Gal4 (C2), *Gr59d*-Gal4 (C1, C2, C4), *Gr66a*-Gal4 (C1, C2, C3, C4), *Gr59e*-Gal4 (C5), *Gr21a*-Gal4 (C6), *GMR57BO4*-Gal4 (C7). C. UAS-*myrGFP* reporter was expressed in individual GSNs using corresponding Gal4 lines (B). We observed a relatively stereotypic position of specific neurons within the organ across animals (n ≥ 3) – exemplified on C6 and C1. D. Illustrative 3D map of TOG segmented cells. Position of neurons with known identities (white dots) is approximated based on separate immunostainings. E. Representative recording of the larval primary taste organ stimulated with citric acid 100mM. Cytoplasmic expression of *GCaMP6m* and nuclear expression of *RFP* in all neurons (upper left panel) and cell segmentation (white spots, lower left panel). Responding neurons are indicated by blue, red and magenta arrows (upper right panel) and by dots and corresponding fluorescence traces of same colors (lower right panels).

**Figure 2:**
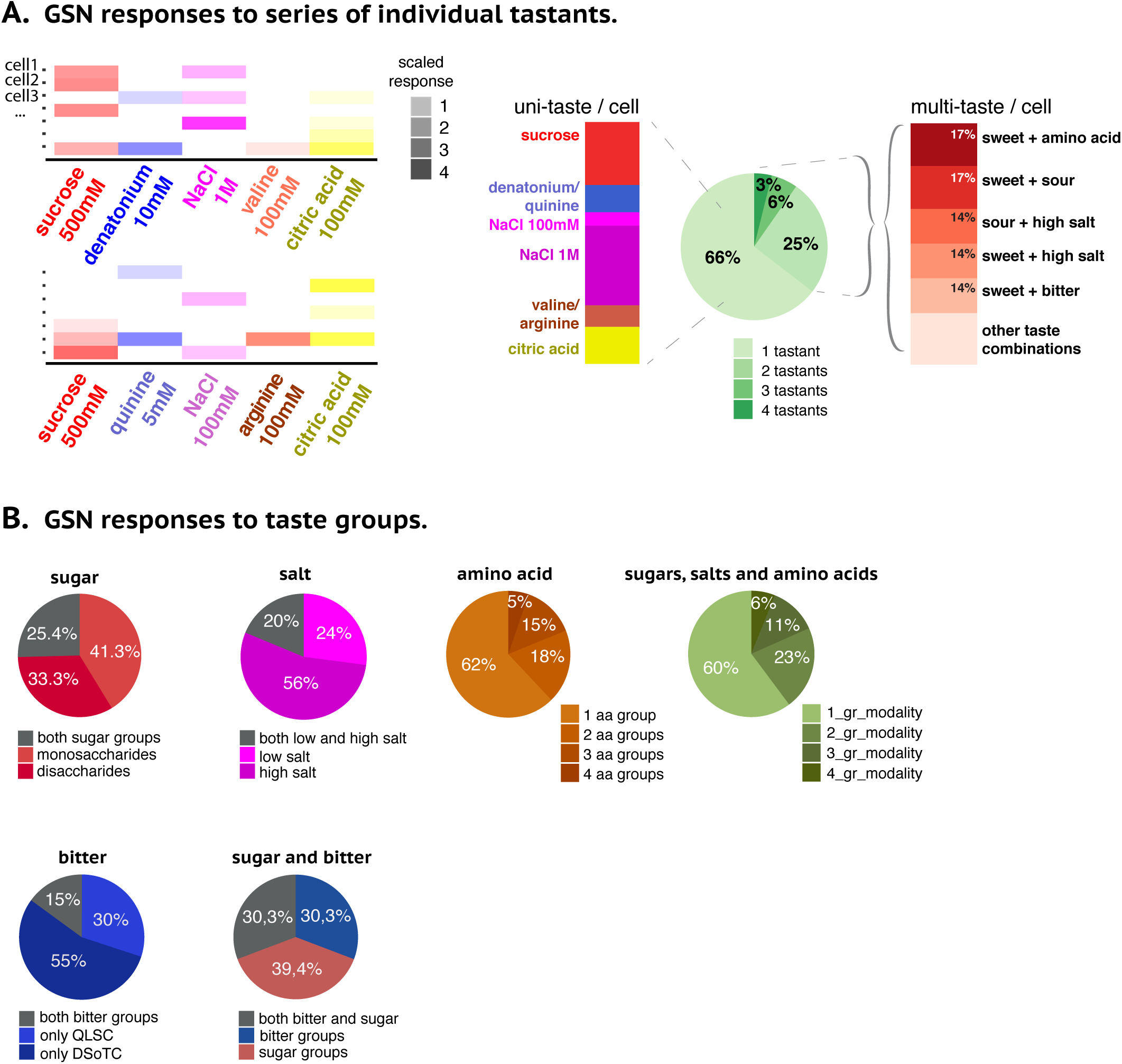
Taste neurons show high proportions of taste multimodality and opposed valence integration. A. GSN responses to series of individual tastants. Illustrative cell responses (y axis) for each tastant series (x axis). Colors were scaled to reflect the intensity of activation to the given taste. Only responses above 20% DF/F were taken into account. Taste stimulation was done with 5 tastants per animal, each belonging to one of the 5 canonical taste modalities represented along the x axis: sweet (shades of red), bitter (blue), salt (purple), amino acids (brown) and sour (yellow). We used 2 different series of taste recordings. Series 1 consisted of sucrose 500mM for sweet taste, denatonium 10mM for bitter, NaCl 1M for high salt, valine 100mM as amino acid and citric acid 100mM for sour taste; correspondingly, the second series consisted of sucrose 500mM, quinine 5mM, Nacl 100mM for low salt, arginine 100mM and citric acid 100mM. 7 separate organs were recorded within the first series and 8 within the second (N=15). Total percentages of taste integration per neuron per animal are represented on the pie chart: 66% of total responding neurons were activated by only one tastant per organ and up to 34% of neurons responded to more than one tastant. Most unimodal responses (uni-taste / cell, left bar-chart) were recorded to sucrose and to high salt. Conversely, different combinations of taste categories were observed to activate single neurons, with different frequency of occurrence (multi-taste / cell, right bar-chart). The most frequent combinations of taste modalities per cell per animal were sweet + amino acid, sweet + sour, sour + high salt, sweet + high salt and sweet + bitter. Some of these combinations involve a presumed positive valence (sucrose/sweet) taste together with a negative valence taste (bitter or high salt) sensed by the same neuron. B. Stimulation with groups of tastants comprising 2 sugar categories (mono and disaccharides), low and high salt concentration, 4 amino acid and 2 bitter groups. Responses of above 20% DF/F intensity threshold were considered. Upper panels: Among all neurons responding to sugar group stimulation, we observed a comparable ratio of cells activated by only one sugar category and 25.4% activated by both mono and disaccharides. For salt-responding neurons, a higher proportion showed unique activation to high salt than to low salt concentration, and 20% neurons responded to both categories. Amino acids split in 4 groups gave a rounded 62% to 38% ratio of uni-to multi-group response integration, calculated among all amino acid-responding neurons. Taking into account total responding neurons from animals stimulated with all 8 groups resulted in 60% to 40% uni-to multi-modality group response integration. Lower panels: Within the total number of bitter-responsive neurons, 55% were uniquely activated by stimulation with DSoTC group and respectively 30% by QLSC group. Integration between sweet and bitter taste has been calculated on responding neurons from animals stimulated with 2 groups of each modality: we observed similar proportions of taste cells responding only to bitter groups (30,3%), cells responding only to sugar (39,4%) and cells responding to groups from both modalities (30,3%).

Although deactivation responses were noted to different substances in different neurons (Fig.4A, blue ovals), such events with frequency of response above 20% were only mapped to CDL2 and CPV neurons, particularly for citric acid stimulation. Noteworthy, CDL1 was robustly activated by citric acid while its CDL2 neighbor showed robust deactivation in response to citric acid (Suppl.File2, Wilcoxon signed rank nonparametric test, p<0.05) (Fig.4C), as described for their concomitant response pattern in figure 3.

**Figure 3:**
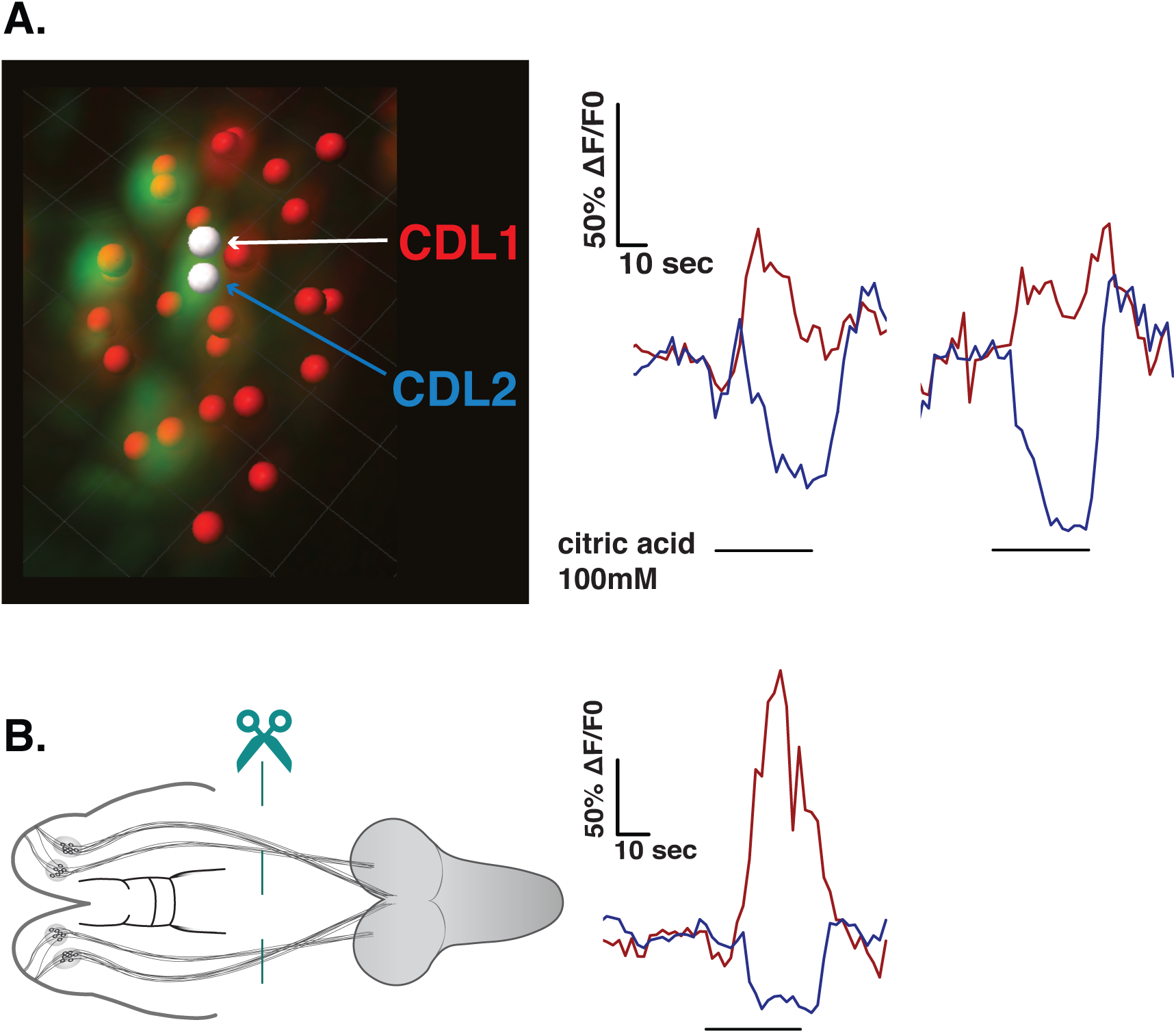
GSNs can respond to taste through activation or deactivation. A. Illustration of a segmented TOG with two highlighted neighboring neurons (white dots) called CDL1, labeled in red and corresponding red fluorescence traces, and CDL2 with corresponding blue traces. When stimulated with citric acid 100mM CDL1 generally responds with a canonical activation of fluorescence intensity increase, while the adjacent CDL2 responds with a simultaneous deactivation, or a fluorescence intensity decrease. B. Cutting connections between chemosensory organs and brain (left panel illustration) did not abolish the synchronous activation-deactivation response to citric acid stimulation observed in CDL1-CDL2 neuronal couple (right panel).

**Figure 4:**
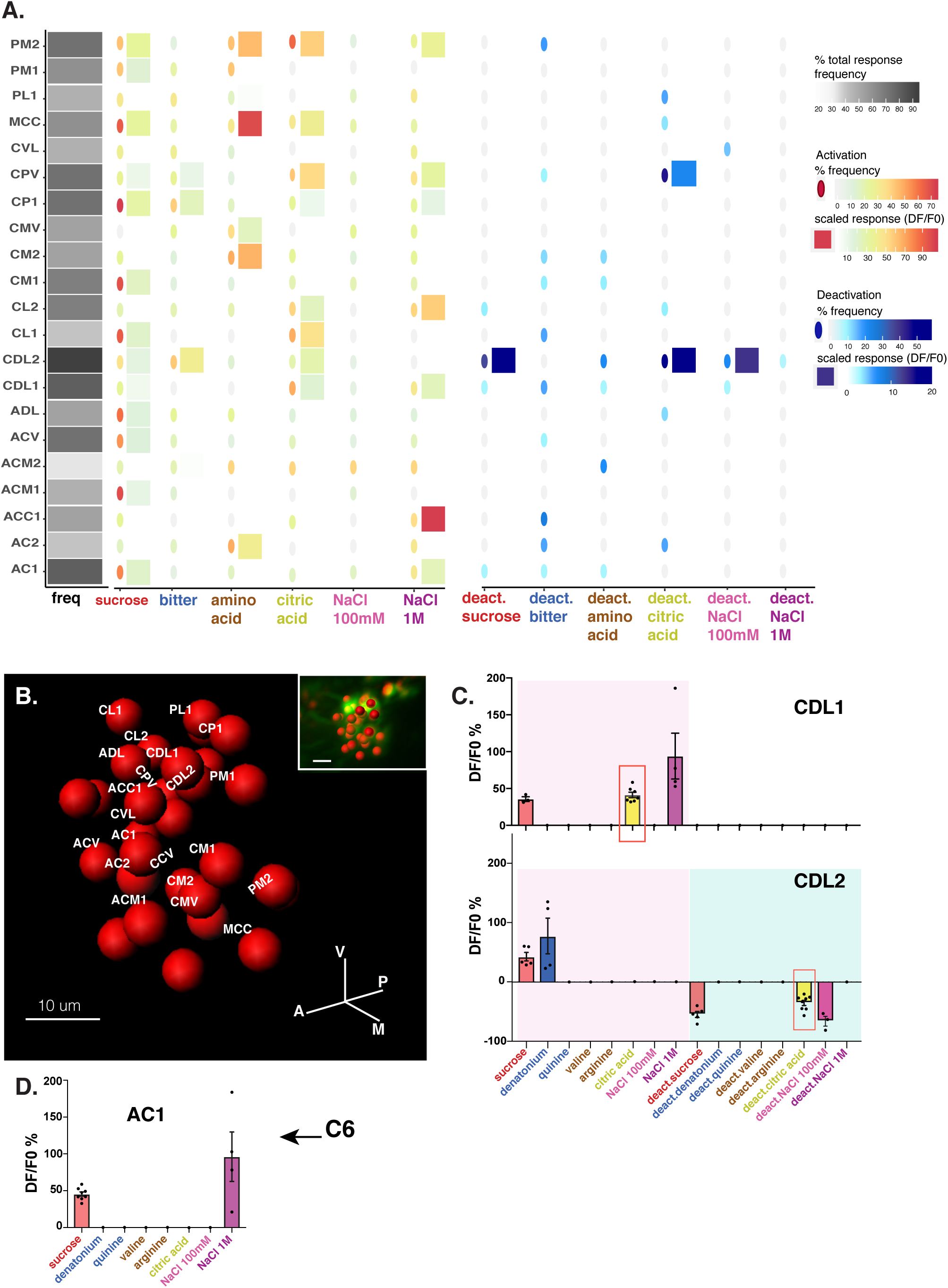
Mapping neuronal responses shows a high level of taste multimodality. A. Manual matching of responding GSNs across recorded organs (N=15) stimulated with series of individual tastants (Fig.2A). 21 neuronal identities displayed on the y-axis. The total response frequency (freq) of each neuron is represented in shades of grey (darker for higher frequency) and translates how often the given neuron has been recorded to respond above 20% DF/F0. For each mapped neuron the activation (yellow-red colors) and deactivation (light-dark blue colors) to a respective tastant is displayed as heatmap for both frequency and amplitude of response. Frequency of response (ovals) shows how consistently a mapped neuron responded to a particular tastant. Response amplitudes (squares) were plotted only for frequencies above 20% (N≥3), scaled and reflected by color intensity of squares. To ease overall interpretation, denatonium and quinine were grouped under the label “bitter”, and valine and arginine as “amino acid”. B. 3D representative visualization of matched neurons. C. CDL1 is activated by sucrose, citric acid and high salt, responding most robustly (most frequently) to citric acid (red square, upper panel). By contrast, CDL2 is activated by sucrose and denatonium, eliciting fluorescence deactivation to sucrose, low salt and most frequently to citric acid (red square, lower panel). D. AC1 is potentially the previously characterized C6 based on the very particular anterior position in the organ (as shown in Fig.1C) and on detected responses to sucrose and high salt, as previously described (48).

While most GSN identities cannot be pinned down with absolute certainty, we identified the AC1 neuron to likely share identity with the previously described C6 neuron, based on the easily recognizable anterior position in the organ (Fig.1C & 4B) and response profiles to sucrose and high salt as previously reported for C6 (48) (Fig.4D).

Taken together, the physiological results show an important degree of taste integration and complexity – a tastant can elicit responses from multiple GSNs and a GSN can respond to multiple tastants per animal or across animals, suggesting a combinatorial mode of sensory encoding.

### Molecular profiles of TOG and DOG neurons at single cell resolution

In order to assess molecular features of individual sense neurons we performed single cell transcriptomic experiments on the TOG and the DOG. Due to low neuronal number and to restricted access to the tissue, the choice of methodology was a limiting and important factor. To generate single cells in suspension at the level of sense organs we proceeded with their isolation by manual picking (Fig.5A) and importantly, we employed a deterministic DropSeq approach for low cell numbers (50, 61), resulting in increased efficiency of encapsulation compared to classical approaches (62, 63).

**Figure 5:**
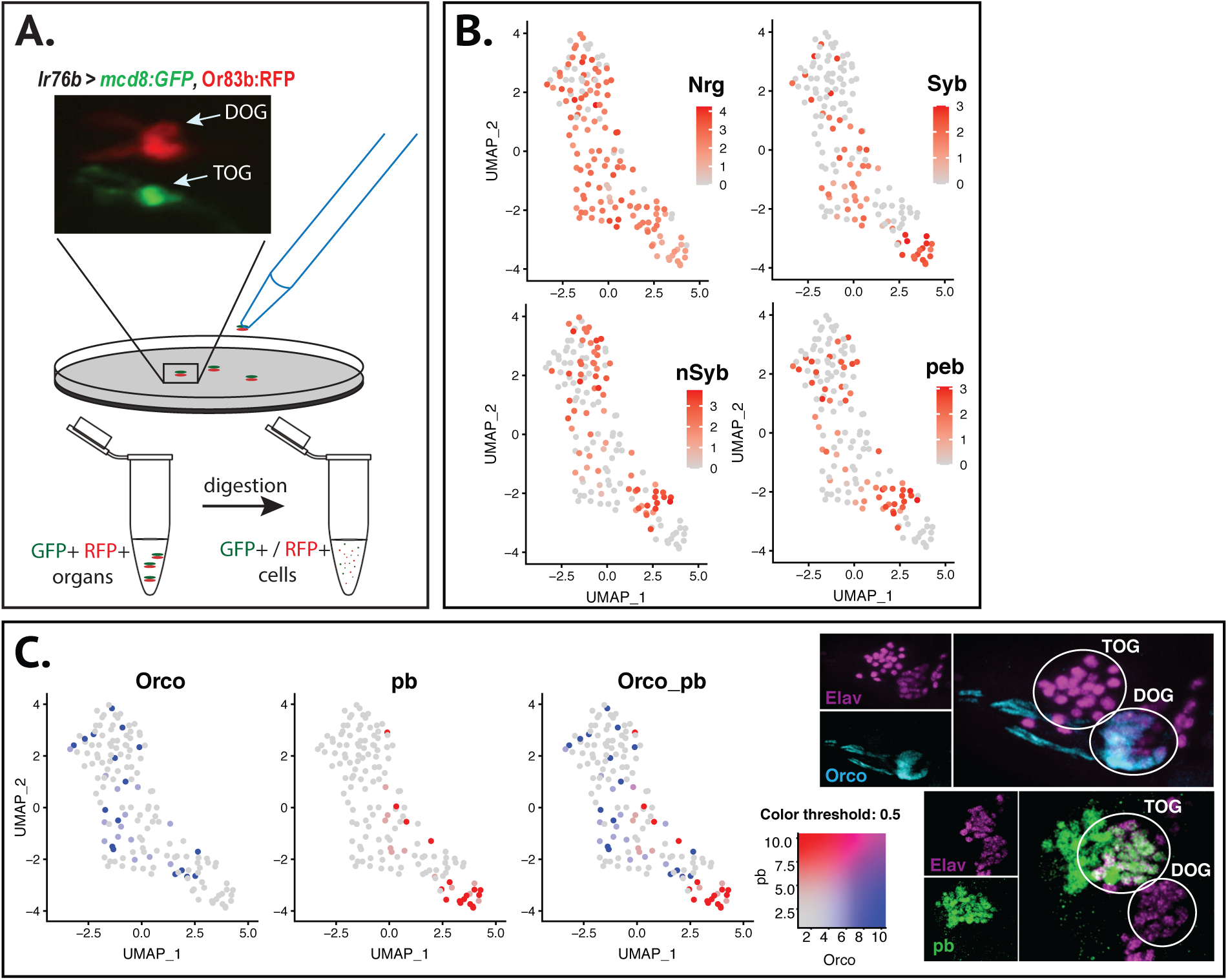
Single-cell RNA sequencing of chemosensory neurons. A. Single cell dissociation protocol; manual picking of the fluorescently labelled sense organs. For chemosensory organ identification, green fluorescence can be driven in all neurons by a pan-neuronal driver line or in a subpopulation as labeled by *Ir76b. RFP* as a tag to *Orco*, olfactory co-receptor present in olfactory cells, labels only DOG. B. Integrated data sets as a single object in Seurat, represented in an UMAP embedding. The final dataset after processing contains 153 cells expressing neuronal markers such as *Neuroglian (Nrg), Synaptobrevin (Syb), neuronal Synaptobrevin (nSyb)* or *pebbled (peb)*. C. *Odorant receptor co-receptor* (*Orco)* is expressed in the DOG and *proboscipedia (pb)* in the TOG. Their molecular expression appears largely non-overlapping in single cells. Immunostainings against the RFP-tagged *Orco* showed expression restricted to the DOG, as expected; *pb* expression shown by anti-Pb staining at embryonic stage labels TOG but not DOG neurons.

The single cell data was analyzed in Seurat v3.1.2 (64, 65) (Suppl.Fig.5B, see Methods), obtaining a dataset of 153 cells with more than 400 genes per cell. We used UMAP embedding for visualization of marker gene expression (Suppl.Fig5C) and confirmed neuronal identity of the relatively homogenous cell population based on neuronal gene markers (Fig.5B). Moreover, we could detect distinct and non-overlapping expression of marker genes for gustatory and olfactory sensory cells. Specifically, the *Orco* gene (*Odorant receptor co-receptor*) that is present in all olfactory neurons in the DOG (43), most probably labels the olfactory neuronal population. Conversely, *proboscipedia (pb)*, a gene known to mediate specification of adult mouthparts (66), shows no overlap of expression with *Orco*- positive cells, labeling a distinct neuronal population (Fig.5C, left). Importantly, immunofluorescence staining in embryos shows *pb* expression restricted to TOG neurons and non-overlapping with *Orco* (Fig.5C, right). Furthermore, expression of the two marker genes largely correlated with the two generated clusters, although less defined for *Orco*, which shows a more dispersed cell expression pattern (Suppl.Fig5C).

### Chemosensory neurons display broad and narrow expression of neurotransmitter markers

To further explore molecular neurotransmitter profiles of sense neurons we performed immunostainings of marker genes on distinct neuromodulator pathways. Using *Cha*-Gal4 (67), we found *Choline acetyltransferase (ChAT/Cha*) in virtually all sense neurons in imunofluorescence staining (Fig.6A, left), corresponding to a relatively broad molecular expression (Fig.6A, right). Conversely, we identified *pale (ple)* as being expressed in 2 cells of the TOG as shown by TH antibody staining (Fig.6B, left), and corresponding to a cell group identified in the UMAP embedding colored by expression (Fig.6B, right). *Pale* encodes a tyrosine hydroxylase as the first step on dopamine synthesis pathway and is orthologous to human *tyrosine hydroxylase (TH)*.

**Figure 6:**
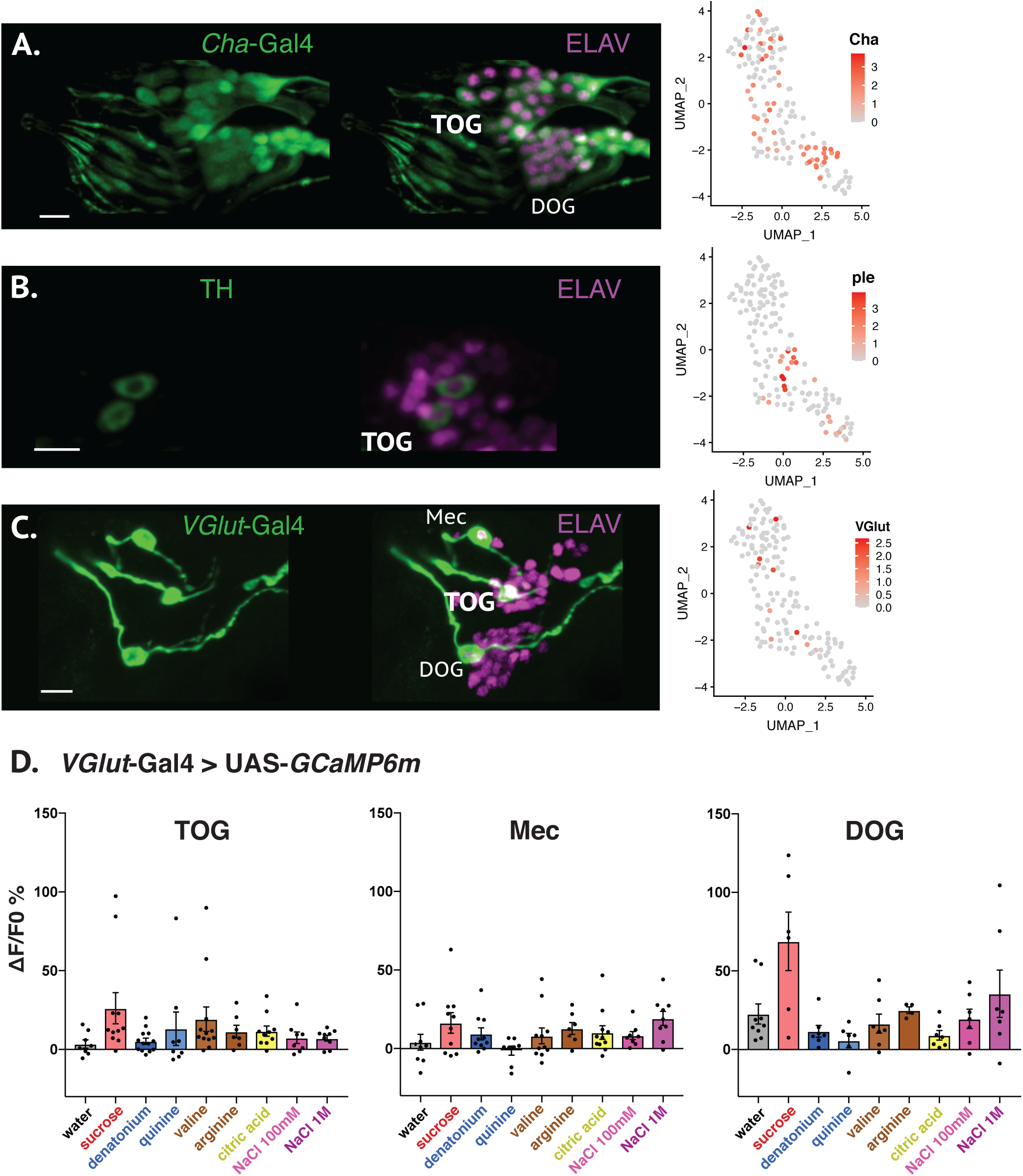
Neurotransmitters of chemosensory neurons show broad or narrow expression. A. *Choline acetyltransferase* (*ChAT*/*Cha*) labels all chemosensory neurons using a *Cha*-Gal4 driver (67) (left panels). Corresponding expression in UMAP embedding colored by expression (right panel). Scale bar, 10 µm. B. *TH* or *pale* (*ple*) is a tyrosine hydroxylase on the dopamine synthesis pathway, expressed in 2 TOG neurons as shown by anti-TH antibody staining (left panels). Corresponding *ple* molecular expression (right panel). Scale bar, 10 µm. C. *Vesicular glutamate transporter* (*VGlut*) shows restricted expression in 3 neurons, using a *VGlut*-Gal4 line (left panels). *VGlut* narrow expression reflected in the UMAP embedding (right panel). Scale bar, 10 µm. D. Physiological responses of the 3 *VGlut* expressing cells stimulated with water as control and with eight tastants introduced earlier (Fig.2A & 4). Mann-Whitney statistical test was used for comparisons between responses to water and each tastant (Suppl.File3): TOG neuron (left panel) responded to sucrose and to valine; Mec (neuron localized anterior-laterally to TOG, middle panel) showed no responses distinct from water; DOG neuron (right panel) responded to sucrose. Data presented as mean + SEM (error bars) and dots represent individual responses from separate experiments.

Other neurotransmitter markers have also shown selective molecular expression, as in the case of *Vesicular glutamate transporter* (*VGlut)*. Immunofluorescence on *VGlut*-Gal4 larvae exposes three cells – one in the TOG, one in the DOG and a third cell in a group peripheral to TOG, we named here Mec (Fig.6C). We verified the response profile of these neurons by recording their physiological response to tastants introduced earlier (Fig.2A), and we compared the response for each tastant with the water response. We noted sucrose responses to be different than control in the TOG and the DOG neurons (Fig.6D, Suppl.File3, p<0.05, Mann-Whitney test), whereas Mec cell didn’t show responses distinct from water to any of the tested substances. The TOG *VGlut*-positive cell did not show co-expression with C7, C2 or C1-Gal4 (Suppl.Fig6), its identity remaining to be determined.

We have detected co-expression of markers on shared neuromodulatory pathway, as in the case of *Dopamine transporter (DAT)* and *Vesicular monoamine transporter (Vmat)* (Fig.7A). Considering that *Cha* is expressed in all chemosensory neurons according to immunofluorescence observations (Fig.6A), co-expression could be theoretically deduced between this and any other neurotransmitter revealed by the molecular data. Specifically, we have detected *Cha* and *DAT* co-localization in a small subset of cells as illustrated in Fig.7B. Additionally, we have identified several neurotransmitter receptors linked to GABA, serotonin, octopamine or dopamine signaling pathways, some of them showing co-expression in certain neurons, as for example *GABA-B-R2* and the serotonergic receptor *5-HT2B* (Fig.7C). Moreover, neuropeptide expression was also abundant in our data, such as neuropeptide-like 2/4 precursors (*Nplp2, Nplp4*), also co-expressed with other neurotransmitter markers such as *VGlut* or *Vmat* (Fig.7D & E). Co-expression of several neuromodulators in numerically restricted neuronal tissues such as the TOG and DOG supports the multimodal character of these sense organs but the functional implication of this observation remains an open question.

**Figure 7:**
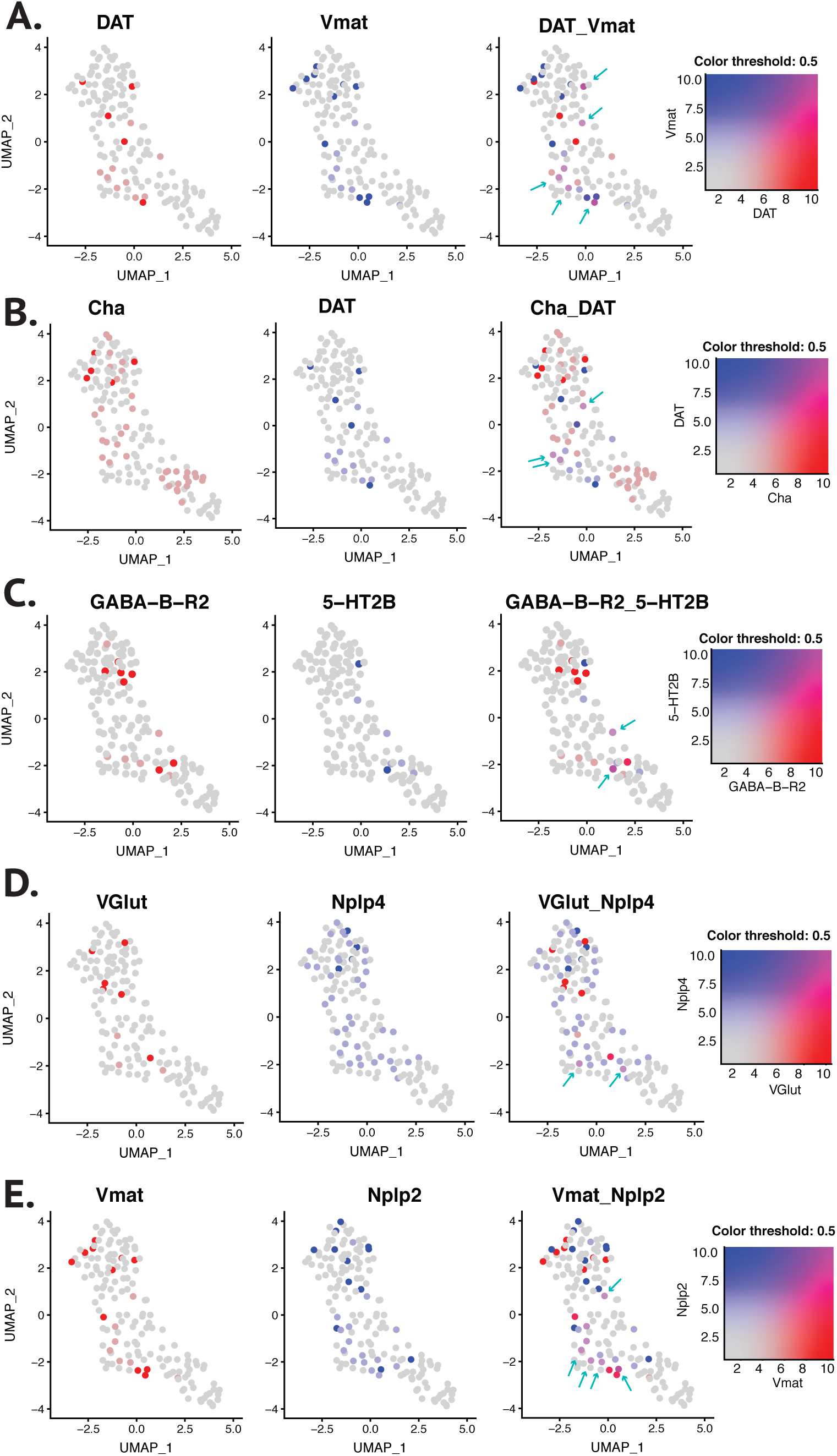
Co-expression of neurotransmitter markers. A. *DAT* and *Vmat* share expression in single cells (magenta dots, blue arrows), as transporters of dopamine and respectively monoamines, the later inclusive of the first. B. Double cholinergic and dopaminergic profile as illustrated by a few cells (blue arrows). C. Co-expression of GABAergic *GABA-B-R2* and serotonergic *5-HT2B* receptors. D. Neuropeptide *Nplp4* present in a few *VGlut*-positive cells. E. Neuropeptide *Nplp2* co-localizes with *Vmat* (blue arrows).

### GSNs show mechanosensory marker expression and responses

Mechanosensory function in the larval taste sensing was assumed to a large extent due to early microscopic descriptions of chemosensory organs in *Drosophila* and in other insects (9, 68, 69). Although functional investigations of mechanical and chemosensory integration have been explored in the adult fly (70, 71), similar studies in the larva were primarily focused on food texture effects in taste behavior and learning (72, 73) while the involved mechanoreceptors remain unknown. We identified expression of mechanosensory markers both by means of immunofluorescence stainings and in single cell UMAP embedding (Fig.8). Namely, *nanchung* (*nan*) is expressed in one TOG and one DOG cell (Fig.8A), whereas *no mechanoreceptor potential C* (*nompC)* is present in a group of 3-4 cells of the TOG and surrounding smaller sensilla (Fig.8B). We found expression of *painless* (*pain*-Gal4) by means of immunolabelling in a population of up to 12 TOG cells, corresponding to a few single cells in the UMAP embedding. Knowing that *pain* is involved in mechanical and thermal nociception in multidendritic and chordotonal sensory neurons in the larva (74), we further tested the response profiles in *pain*-expressing TOG neurons to mechanical stimulation (Fig.8C).

**Figure 8:**
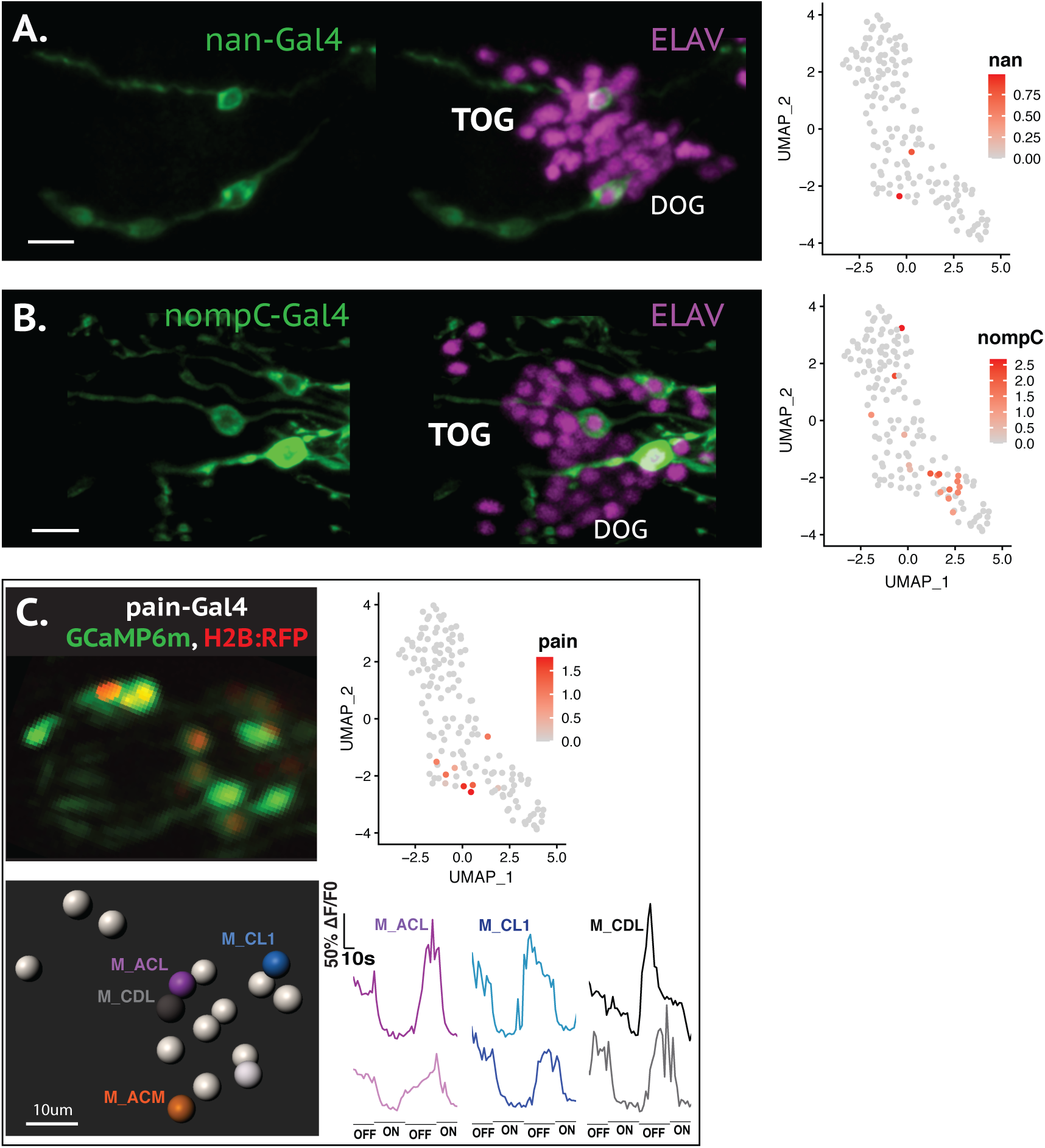
Chemosensory neurons express mechanosensory markers. A. *nan-*Gal4 labels 1 TOG and 1 DOG neuron (left panel), scarce expression mirrored by the UMAP embedding colored by expression (right panel). Scale bar, 10 µm. B. *nompC-*Gal4 has been identified in 3 chemosensory cells (left panel), paralleled by UMAP embedding (right panel). Scale bar, 10 µm. C. *pain*-Gal4 labels a larger subpopulation of TOG neurons. We recorded neuronal response of *pain*-expressing neurons pre-exposed to water: mechanical stimulation produced by water pump switch ON/OFF. Responding cells in a 3D representation and respective fluorescence traces. Representative traces for 3 cells, each from two different animals.

To measure mechanical responses we aimed at inducing fluid shear stress on the cellular membrane (75) and measuring the cell response after turning on the water flow inside the microfluidic chip. As the tissue was exposed to water also during the off stimulus (in absence of flow), we interpreted neuronal responses to be determined by the fluid flow and not by water taste itself. We detected fluorescence intensity changes on average in 4 *pain*-Gal4 cells, which were named based on their relative position and mapped to the organ. More often than not, although not exclusively, we observed fluorescence intensity decrease during the on stimulus, *pain*-expressing cells being potentially de-activated by shear stress (Fig.8C).

In a separate assay we recorded *nsyb*-Gal4 larvae for mechanosensory as well as taste sensing. Interestingly, in few cases we observed neurons activated by both types of modalities, as for example CM1 neuron responding to 500mM sucrose but also showing de-activation to mechanical shear stress (Suppl.Fig8). This observation suggests that some GSNs could integrate both mechano- and taste sensing as previously suggested (8).

## Discussion

In this study we explored the physiological and molecular characteristics of *Drosophila* larval taste system. We designed an experimental framework allowing calcium imaging in all GSNs comprised by the larval primary taste-sensing organ at the periphery (the TOG, Fig.1, Suppl.Fig.1). Our approach complements previous strategies based on labeling of few single neurons or subpopulations (6, 29, 35, 39, 48), allowing for an overview of activation patterns within the organ. Strengthening our previous observations on the multimodal character of larval taste neurons (48), we identified single GSNs activated by different taste categories as well as single GSNs activated by tastants with opposite associated valence. Cell integration of different tastes might involve spatio-temporal codes, as described for gustatory discrimination in the moth (76) or combined patterns of response in cell number and type of activation. In line with our observations that one tastant elicits response in several GSNs, it is interesting to consider that the relevance of a stimulus and ultimately its behavioral output could depend on neurons firing together, concurrent activation of 2 neurons differing from their separate activation (77).

For the substances tested within this study we observed roughly a 60-70% versus 30-40% division of uni-modal versus multi-modal responding GSNs (Fig.2 & 4), comparable with tuning proportions reported for mouse fungiform taste sensing cells (78). The *best stimulus concentration* principle is arguably the strongest support of the labeled-line taste coding at the periphery in the mammalian system (79). However, this model leaves out the case of tastants activating different types of taste receptors depending on concentration as in the case of acesulfame K and saccharin, which at low concentrations activate sweet receptors but with increased concentration start activating bitter receptors (12). Accordingly, the chosen substance concentrations within our assay might cause variability in response profiles, and doubtlessly, modifying the dilution for the same tastants would alter the observed tuning of GSNs, as shown on afferent taste neurons in mammals (15).

An intriguing aspect of taste physiology revealed by our observations was GCaMP fluorescence decrease in GSNs, termed here *deactivation* for ease of discussion. Most notable deactivation signals were recorded in CDL2 neuron and most stereotypically to citric acid (Fig.3 & 4), although the phenomenon was not restricted to this neuron or tastant. Interestingly, citric acid elicited concomitant activation in the neighboring cell named CDL1, indicating response patterns of activation-deactivation in the CDL1-CDL2 couple, similar to ephaptic signaling identified in the olfactory system of the adult fly (51). The synchronous responses were not abolished when severing afferent neuronal axons (Fig.3B), implying that CDL2 deactivation is not determined by a top-down inhibition from brain interneurons. Additional possible explanations of the observed neuronal activity include gap junction communication or tonic activity in CDL2 inhibited by citric acid stimulation, as reported in sensory neurons of bumblebees and respectively in *C*.*elegans* (81, 82). However, the observed synchronous activation coupled with deactivation responses in adjacent neurons best fit the description of ephaptic signaling (60, 80), pointing towards inter-cellular communication and thus emphasizing a model of signal integration in larval taste system at the periphery.

Using DisCo allowed us to successfully probe larval chemosensory organs with single cell resolution (Fig.5, Suppl.Fig.5), resolving low cell input and tissue accessibility limitations. We identified *pb* and *Orco* as marker genes distinctly expressed in TOG and respectively in DOG neurons. If *Orco* is an olfactory co-receptor expressed in all olfactory neurons, taste neuronal population is less characterized and functionally more diverse. Therefore, *pb* could be an interesting candidate for further characterizing the population of *pb*-expressing GSNs.

Moreover, in the present work we describe distinct expression patterns of neuromodulator marker genes in TOG and DOG neurons, as shown by UMAP embedding and confirmed by immunostainings (Fig.6). The cholinergic profile of larval chemosensory neurons has previously been reported (31, 53) but we show that some cells express glutamatergic, dopaminergic and neuropeptidergic markers. Interestingly, marker genes on different signaling pathways are co-expressed in subsets of cells (Fig.7), as previously demonstrated in *Drosophila* CNS (54, 83). Considering the multimodal character of some larval GSNs, it would be fascinating to uncover the role played by co-expression of distinct neurotransmitter markers in such neurons. Nevertheless, if remains surprising that a handful of sensory neurons display a diverse neurotransmitter profile, as reported here, demonstrating a large molecular diversity within the neuronal sensory population.

Furthermore, we found expression of mechanosensory markers as revealed by UMAP embedding of single-cell data and verified with immunostainings. We detected *nan* in one TOG neuron and one DOG neuron, whereas *nompC* and *pain* label a larger population. We recorded responses of *pain*–expressing GSNs exposed to water flow turned on and off, as we intended to create fluid shear stress on the cell membrane, similarly to observations on cultured cells (84) and studies on vascular tissue (85). It remains to be elucidated if shear stress responses detected in *pain* neurons are mediated by Painless channel itself. Painless is a protein known to be important in mechanical and thermal nociception in larvae (74), although its role remains unexplored in the gustatory system. On the other hand, Painless has also been shown to be involved in wasabi taste detection in adult flies in Gr66a-expressing GSNs (86) and we have also found scarce co-expression between *pain* and *Gr66a* (data not shown). The presence of mechanosensory neurons in the larval primary taste center has been frequently reported (9, 49). Although chemosensory sensilla have been classically described as distinct from mechanosensory sensilla (87), existence of neurons with dual chemo- and mechanosensory modalities has also been implied based on their tubular dendritic body (8). In our observations, some mechano-responsive neurons also show activation to gustatory cues (Suppl.Fig.8), stressing on the probability of multisensory integration of taste and mechanical cues.

Taken together, our data suggest a complex taste sensory coding in the larva, made up by patterns of activity across neurons rather than by labeled individual neuronal responses for discrimination between tastants. Besides canonical fluorescence increase responses we have depicted fluorescence decrease signals in specific neurons pointing towards electric coupling and therefore sensory cue processing at the periphery. Additionally, the neurotransmitter and mechanoreceptor expression contribute to the multifaceted chemosensory system to potentially explain the multimodality and multisensory character of specific taste neurons in future studies.

## Methods

### Fly stocks

We used the following fly strains: *nSyb*-Gal4 (gift from Shanhaz Lone), UAS-*GCaMP6m* (Bloomington #42748), UAS-*H2B*:*RFP* (gift from Boris Egger), *GMR57BO4*-Gal4 (Bl. #46355), *Gr94a*-gal4 (Bl. #57686), *Gr21a*-Gal4 (Bl. #23890), *Gr22e*-Gal4 (Bl. #57608), *Gr66a*-Gal4 (Bl. #57670), *Gr97a*-Gal4 (Bl. #57687), *Gr59e*-Gal4 (Bl. #57655), UAS:*mcd8:GFP;Or83b:RFP* (gift from Travis Carney), UAS-*myr:GFP* (Bl. #32198), *VGlut*- Gal4 (Bl. #24635), *VGAT*-Gal4 (Bl. #58980), *Cha*-Gal4 (kindly gifted by PM Salvaterra), *nan*-Gal4 (Bl. #24903), *nompC*-Gal4 (Bl. #36361), *pain*-Gal4 (Bl. #27894), FlyFos *Orco*- GFP (VDRC 318654).

### Immunostainings

3^rd^ instar larvae (4 days after egg laying) were washed and dissected in ice-cold PBS. Tissue containing chemosensory neurons was fixed in 3,7% formaldehyde for 18-20min at room temperature. After fixation, incubation in primary antibodies was performed at 4°C overnight, then in secondary antibodies at 4°C overnight, and finally, in Vectashield antifade medium (Vector Laboratories) for at least 1h before mounting the samples on microscope slides. After each step described above, PBST (PBS 0,3% Triton X-100) was employed for three consecutive plus three 30-minute washes at room temperature.

Immunolabelling against Pb (Fig.5C) was performed at embryonic stage. Embryos were washed in tap water and placed in a plastic mesh in a sampling manifold. (Millipore XX2702550). First, they were dechorionated in 2.5% sodium hypochlorite (household bleach) for 2 minutes. Embryos were then fixed in a 1:1 solution of n-heptane:PEM buffer with 3.7% formaldehyde for 20 minutes at room temperature on a rotating wheel. Upon fixation, embryos were washed with methanol and blocked for 1 hour at room temperature with 5% normal goat serum in PBST. The samples were incubated with antibodies and mounted as described above.

Confocal images were acquired on a Leica TCS SPE-5 confocal, using the 40x oil immersion objective for larval staining and the 63X glycerol immersion for embryo staining, at 0.8-1um slice thickness. Images were assembled using Fiji (88) and Adobe Illustrator.

Primary antibodies: chicken anti-GFP (1:1000, Abcam, ab13970), rabbit anti-DsRed (1:1000, Clontech, No. 632496), rat anti-ELAV (1:30, No. 7E8A10), anti-TH (1:100, Millipore, ab_390204), rabbit anti-Pb 1:100 (gift of Thomas Kaufman, Cribbs, 1992 #475). Secondary antibodies conjugated with Alexa Fluor fluorescent proteins (488, 568, 647) were used in dilution of 1:200 (Molecular Probes no. A-11008, A-11039, A-21244, A-21247, A-11011).

### Chemicals

For calcium imaging experiments the highest purity available was chosen for all chemicals used in this study: L-Alanine (Sigma-Aldrich, 56-41-7), L-Glutamine (Sigma-Aldrich, 56-85-9), L-Aspartic acid (Sigma-Aldrich, 56-84-8), L-Glutamic acid (Sigma-Aldrich, 56-86-0), L-Asparagine (Sigma-Aldrich, 70-47-3), L-Proline (Sigma-Aldrich, 147-85-3), L-Tyrosine (Sigma-Aldrich, 60-17-4), L-Threonine (Sigma-Aldrich, 72-19-5), L-Phenylalanine (Sigma-Aldrich, 63-91-2), L-Lysine (Sigma-Aldrich, 56-87-1), L-Arginine (Sigma-Aldrich, 74-79-3), L-Histidine (Sigma-Aldrich, 71-00-1), L-Serine (Sigma-Aldrich, 56-45-1), L-Valine (Sigma-Aldrich, 72-18-4), L-Cysteine (Sigma-Aldrich, 52-90-4), L-Methionine (Sigma-Aldrich, 63-68-3), L-Leucine (Sigma-Aldrich, 61-90-5), L-Isoleucine (Sigma-Aldrich, 73-32-5), L-Tryptophan (Sigma-Aldrich, 73-22-3), L-Glycine (Roth, 3187.3), D-Sucrose (Sigma-Aldrich, 57-50-1), D-Glucose (Sigma-Aldrich, 50-99-7), G-Fructose (Fluka, 57-48-7), D-Trehalose dihydrate (Roth, 9286.1), D-Arabinose (Sigma-Aldrich, 10323-20-3), D-Maltose monohydrate (Sigma-Aldrich, 6363-53-7), D-Mannose (Sigma-Aldrich, 3458-28-4), D-Galactose (Roth, 4979.1), D-Cellobiose (Roth, 6840.3), Lactose monohydrate (Roth, 8921.1), Quinine hemisulfate salt monohydrate (Sigma-Aldrich, 6119-70-6), Denatonium benzoate (Aldrich, 3734-33-6), Caffeine anhydrous (Fluka, 58-08-2), Coumarin (Sigma-Aldrich, 91-64-5), Sucrose octaacetate (Aldrich, 126-17-7), Lobeline hydrochloride (Aldrich, 134-63-4), Theophyline (Sigma-Aldrich, 58-55-9), Strychnine (Roth, 4843.1), Potassium chloride (Merck, 7447-40-7), Sodium chloride (Sigma-Aldrich, 7647-14-5), Citric acid (Sigma-Aldrich, 77-92-9).

### Chemicals used in cell dissociation experiments

elastase (Sigma-Aldrich, E0258), collagenase (Sigma-Aldrich, C9722), PBS (Thermo Fisher Scientific, AM9624), BSA (Thermo Fisher Scientific, AM2616), Murine inhibitor (New England Biolabs (NEB, M0314L).

### Calcium imaging

The semi-intact sample for calcium-imaging recordings was prepared as previously described (57). 3^rd^ instar larval heads comprising chemosensory neurons, the brain and the connecting nerves were dissected in AHL (adult hemolymph like) saline solution. The tissue was introduced into the microfluidic chip and the sample was connected to the tubing and micropump setup. For whole organ TOG recordings UAS-*GCaMP6m* (55) was used to genetically label all neurons and *RFP* was expressed in nuclei using UAS-*H2B:RFP* (56). Stacks were set to 15 slices of 1,5 µm thickness each, pixel binning of 2, with resulting time acquisition speed of 0.5 stacks (time-points) per second. For recordings from *VGlut*-Gal4 neurons, we used the same UAS reporter lines to image the three neurons belonging to the TOG and the DOG, but due to a deeper tissue extending over both organs, we set the stacks to 25 slices of 2,5 µm thickness each, preserving the time acquisition speed.

The microscope of choice was a spinning-disk confocal Visitron CSU-W1. Recordings were exported as .nd files and further processed in Fiji/ImageJ and Imaris software. For each recording an RFP stack was acquired prior to the CGaMP time-series.

### Taste Stimulation

Each tastant was administered using a micropump (mp6, Bartels Mikrotechnik) controlled in VisiView software through built-in macros. Millipore water was used as solvent for taste solutions and as washing control. For each taste stimulation macros were designed for 2-minute recordings to allow 1 minute pre-wash followed by 30-second stimulation and a final 30-second wash.

### Taste stimulation with series of individual tastants

The sequence of chemicals within a trial was randomized. We used two series of tastants (Suppl.Table 1), each containing 5 substances from 5 different taste categories. Series 1 consisted of sucrose (sweet), denatonium (bitter), NaCl 1M (high salt), valine (amino acid) and citric acid (sour). Series 2 consisted of sucrose, quinine (bitter), NaCl 100mM (low salt), arginine (amino acid) and citric acid. Each recorded animal was stimulated with all the tastants within one series. A total of 15 TOG organs were recorded, 7 for series 1 and 8 for series 2.

### Taste stimulation with groups of tastants

We compiled 2 groups of sugars, each containing 5 sugars at a concentration of 100 mM each: monosaccharides (fructose, glucose, arabinose, mannose, galactose), and disaccharides (sucrose, trehalose, maltose, lactose, cellobiose). Amino acids were prepared in 4 groups (A, B, C and D) as previously described by Park et al. (59) (Suppl.Table 1). Tyrosine was included in group B at a concentration of 1mM. Salt taste groups contained NaCl and KCl in equal proportions at a total concentration of 50mM (low salt) and respectively 1M (high salt). Bitter tastants were randomized in 2 groups of 4 substances each, with a final concentration of 22mM and respectively 13mM, restricted by the solubility of certain substances such as lobeline or strychnine reduced to 1mM instead of 10mM (Suppl.Table 1): DSoTC group (denatonium benzoate, sucrose octaacetate, theophylline, coumarin), and respectively QLSC group (quinine, lobeline, strychnine, caffeine).

### Mechanical stimulation

Macros were adjusted to switch water flow between on and off at 30-second interval. The sample was exposed to water in both conditions; therefore, corresponding neuronal responses were interpreted to be result of shear stress mechanical stimulation (75) and not caused by water taste.

### Data processing and analysis of whole-organ physiological recordings

We first carried out deconvolution processing on Huygens platform of RFP stacks and GCaMP recordings. A Fiji/ImageJ plugin was written to duplicate the RFP signal onto all time-points of the CGaMP recording (Suppl.Fig.1A). If the animal movement was considerable, a 3D drift correction was applied on the GFP signal before merging the two channels. Eventual misalignment between the two signals (caused by drifts or animal movement) was corrected by writing a Fiji/ImageJ macro for manually guided realignment (Suppl.Fig.1B). A final 3D drift correction (89) was then performed for better stabilization. The recording was transferred in Imaris for automatic segmentation with spots of 3,5um diameter. Each uniquely identified neuron though spot detection received an identity and a corresponding fluorescence trace over time (Suppl.Fig.1C). An extra spot was created in proximity to the organ, serving as background fluorescence to be subtracted from all cell spots. Signal processing and downstream analysis were performed with R (90, 91). Normalization of fluorescence values followed the formula (F_t_-F0)/F0, F_t_ being the intensity at timepoint t and F0 the baseline as average intensity of 10 frames prior stimulation. Response amplitude was calculated as the difference between the peak intensity after stimulation (average of 5 frames near the maximal fluorescence value) and F0. All traces were individually inspected to exclude false positive responses. Responses were considered for further analysis if their amplitude was minimum 20% DF/F0. To calculate taste integration percentages (Fig.2), for each a specific dataset we selected all responding neurons and then determined the fraction of neurons activated by one or by multiple tastants / taste groups per animal. Illustrative lists of responding neurons as well as number of neurons and responses considered for analysis within each test category were assembled in Supplementary File 1. Neurons responding to individual tastants were mapped across organs (Fig.4) by manually assessing their relative position and physiological response. Supplementary File 2 contains he activation or deactivation values with frequencies of response above 20% for mapped neurons and the corresponding Wilcoxon signed rank nonparametric test p-values. Figures were generated in RStudio (92) using “ggplot2”, “reshape2” and “scales” packages (93-95) or in GraphPad Prism 7 and adjusted in Adobe Illustrator.

### Calcium imaging analysis of *VGlut*-positive cells

For recordings of *VGlut*-Gal4 neurons, analysis was entirely performed in Fiji/ImageJ. Cell segmentation was replaced by setting regions of interest (ROI) around each of the 3 somas and fluorescence intensity over time was extracted after applying a z-projection of the stack recordings. Normalization was performed as above and all taste responses were compared to water response for each neuron. Response values and Mann-Whitney tests of comparisons with control were compiled in Supplementary File 3. Analysis and figures were generated in GraphPad Prism 7.

### Single-cell chemosensory cell suspension preparation for DisCo

3^rd^ instar larvae of genotype *nsyb*-Gal4 > UAS-*mcd8:GFP; Or83b:RFP* were collected from food, washed in tap water, PBS, dropped in ethanol and again PBS and dissected in ice-cold PBS in such a manner that only the external chemosensory organs were kept, avoiding to include also the pharyngeal tissue containing internal chemosensory organs. The isolated material was then placed on ice in elastase 1 mg/ml in siliconized 2-ml tubes. After dissecting 20-30 larvae (20-25 minutes), the tube sample was placed at room temperature to initiate digestion. After 30 minutes the tissues was washed in PBS+BSA0,05% and dissociated by up-down pipetting for 120 times using siliconized 200p pipette tips. Separated TOG and DOG organs were detected using a fluorescence stereomicroscope and manually picked with a glass micropipette, placed in a final dissociation enzyme mix of Collagenase 1mg/ml + Elastase 0,5mg/ml for 10-15 minutes for single cell suspension. The reaction was stopped with PBS + BSA 0,05%. Murine inhibitor was added at each step of the dissociation protocol.

### Deterministic co-encapsulation (DisCo) of chemosensory neurons for single cell transcriptomics

Microfluidic chip design, fabrication and device handling are described elsewhere (50). Following organ dissociation, target cell suspension was diluted in the cell loading buffer containing PBS 0.01 % BSA (Sigma B8667), 6% Optiprep (Sigma D1556) and Murine RNase inhibitor (NEB M0314L) in the loading tip connected to the DisCo chip. After bead-cell in droplet co-encapsulation, sample droplets were transferred to a bead collection chip. Subsequently to bead capture, washing, reverse transcription (Thermo Scientific EP0753) and Exonuclease I (NEB M0293L) reactions were performed on-chip (61). Beads containing cDNA were then eluted and cDNA was amplified for 21 cycles using Kapa HiFi Hot start ready mix (Roche #07958935001). Amplified cDNA was then purified (GC biotech CPCR-0050) for quality assessment with Fragment Analyzer (Agilent). Libraries were then tagmented using in-house Tn5 (96), size selected and purified for sequencing on NextSeq 500 system (Illumina) following recommendations from original Drop-seq protocol (97) (20 bp for read 1 and 50 bp for read2) at sequencing depth above 400.000 reads per cell.

### Single-cell data pre-processing and analysis

The data analysis was performed using the Drop-seq tools package (62). After pre-processing, reads were aligned to *Drosophila melanogaster* reference genome (Ensembl version 86) using STAR (version 2.7.0.e) (98). Following the alignment, BAM files were processed using the initial package and read-count matrices were generated.

Downstream analysis was done using the Seurat package (65) version 3.1.2 in RStudio. Individual data sets were loaded and used to create separate normalized and scaled Seurat objects of minimum 400 genes per cell. In order to apply unique cell filters we merged the data and then excluded cells with high gene number and high UMIs as potential doublets and cells with high mitochondrial gene percentage indicating potential apoptotic cells. Due to observed correlation between cells with high gene number (nGene, Suppl.Fig.5B) and cells with high UMIs (nCount, Suppl.Fig.5B), by applying UMI threshold at 50000 we also eliminated cells with more than 4000 genes. Cells with mitochondrial gene percentage under 9% were kept for further analysis (Suppl.Fig.5B). Data was then integrated to circumvent batch effects using Seurat functions *FindIntegrationAnchors* and *IntegrateData*. As we were interested in characterizing neurons, we used the *subset* function to keep only cells expressing *nSyb* or *peb* neuronal markers, excluding eventual surrounding tissue or cuticle cells. On the final dataset of 153 neurons PCA (principal component analysis) computation was followed by UMAP embedding and clustering (Suppl.Fig.5C) was performed at 0.5 resolution.

## Acknowledgements

We thank the Sprecher lab for support and useful discussions and Joern Pezoldt, Wanze Chen and Maria Litovchenko from Deplancke lab for support, valuable comments and practical advice. We thank EPFL’s Gene Expression Core Facility for sequencing support and Rommelaere Samuel from Lemaitre lab for providing fluorescent microscope during single-cell preparation experiments. We are also grateful to Boris Egger for guidance on microscopy and data analysis and to everyone who shared with us fly stocks and reagents.

## Author contributions

GLM and SGS designed the study and wrote the paper. GLM performed physiological recordings, immunostainings, and single cell dissociation, analyzed the data and created the figures. MB and JB performed DisCo experiments and library preparation. FM wrote the plugins and macros necessary for calcium imaging data processing and assisted in establishing the analysis pipeline. CBA identified and determined *proboscipedia* immunofluorescence expression and assisted with single cell sequencing analysis in Seurat. BD, JYK and SGS assisted in interpretation of results.

## Additional information

The authors declare no competing interests.

## Data Availability

Single cell RNAseq data reported in this study are available on NCBI GEO; accession number GSE149975.

Fiji/ImageJ plugin and macro for calcium imaging data processing are available as supplementary file.

Data generated and analyzed within the present study are available from the corresponding author upon request.

---

**Supplementary Figure 1:**
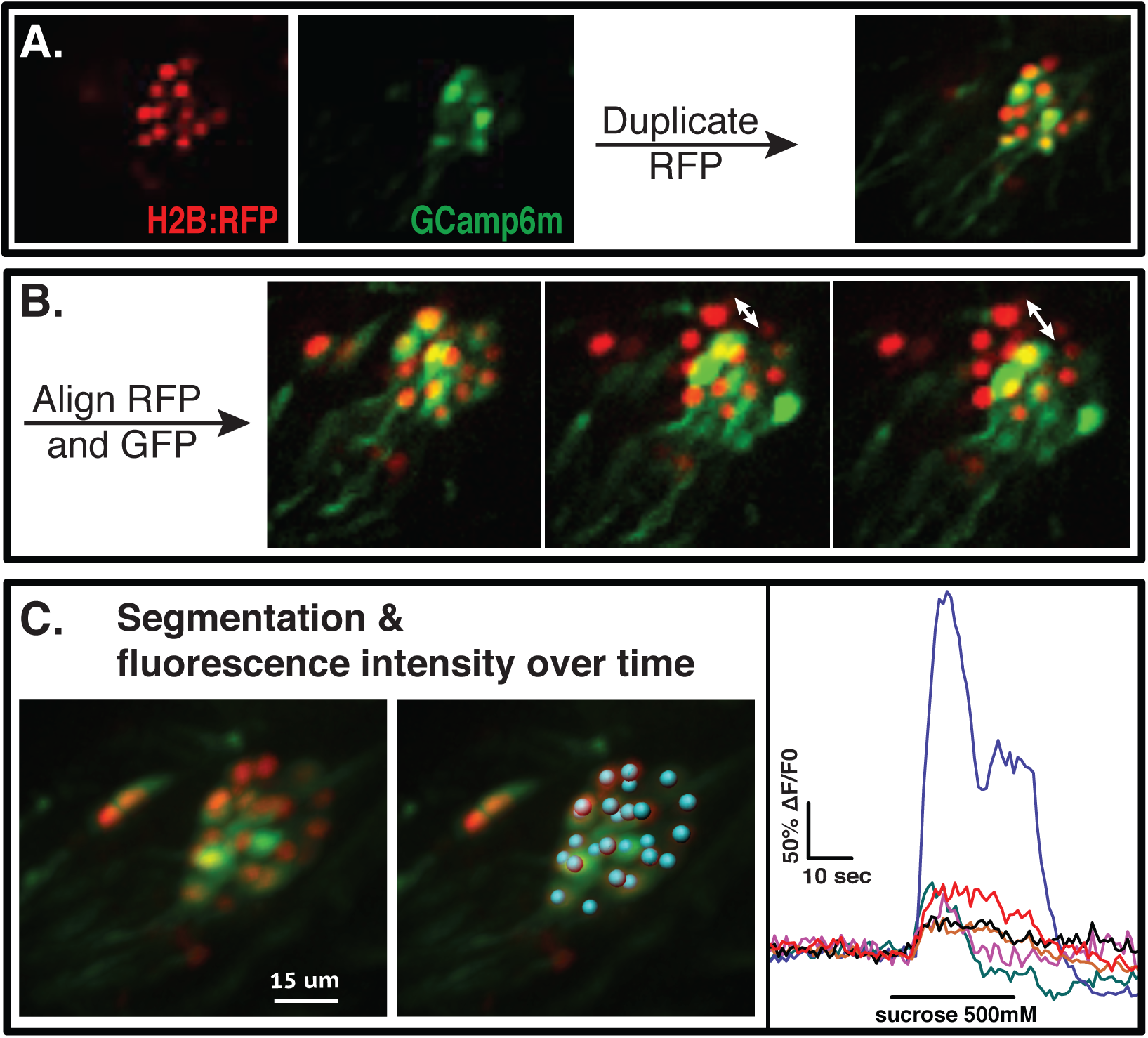
Whole organ calcium imaging – data processing. A. The RFP nuclear signal acquired as a single stack was duplicated to all timepoints of the GCaMP time-series using an in-house written plugin. Step performed in Fiji/ImageJ. B. On the recording comprising both fluorescence channels, any misalignment between the two signals due to animal movement or slight drift was corrected through a manually assisted macro. Step performed in Fiji/ImageJ. C. The resulting stabilized and aligned recording was transferred to Imaris for cell segmentation (middle panel) and extraction of corresponding fluorescence traces (right panel). Traces of neurons responding to sucrose 500mM in a representative recording. Steps performed in Imaris and in RStudio.

**Supplementary Video 1: Whole organ recording and cell segmentation**.

The TOG (larval main peripheral taste organ) recorded using genetically encoded GCaMP reporter in all neurons. The second fluorophore, nuclear RFP, is used for cell segmentation, allowing creation of spots corresponding to each individual neuron. GCaMP fluorescence intensity over time for each spot is represented by spot color, ranging from blue (low signal) to red (high intensity).

**Supplementary Figure 5:**
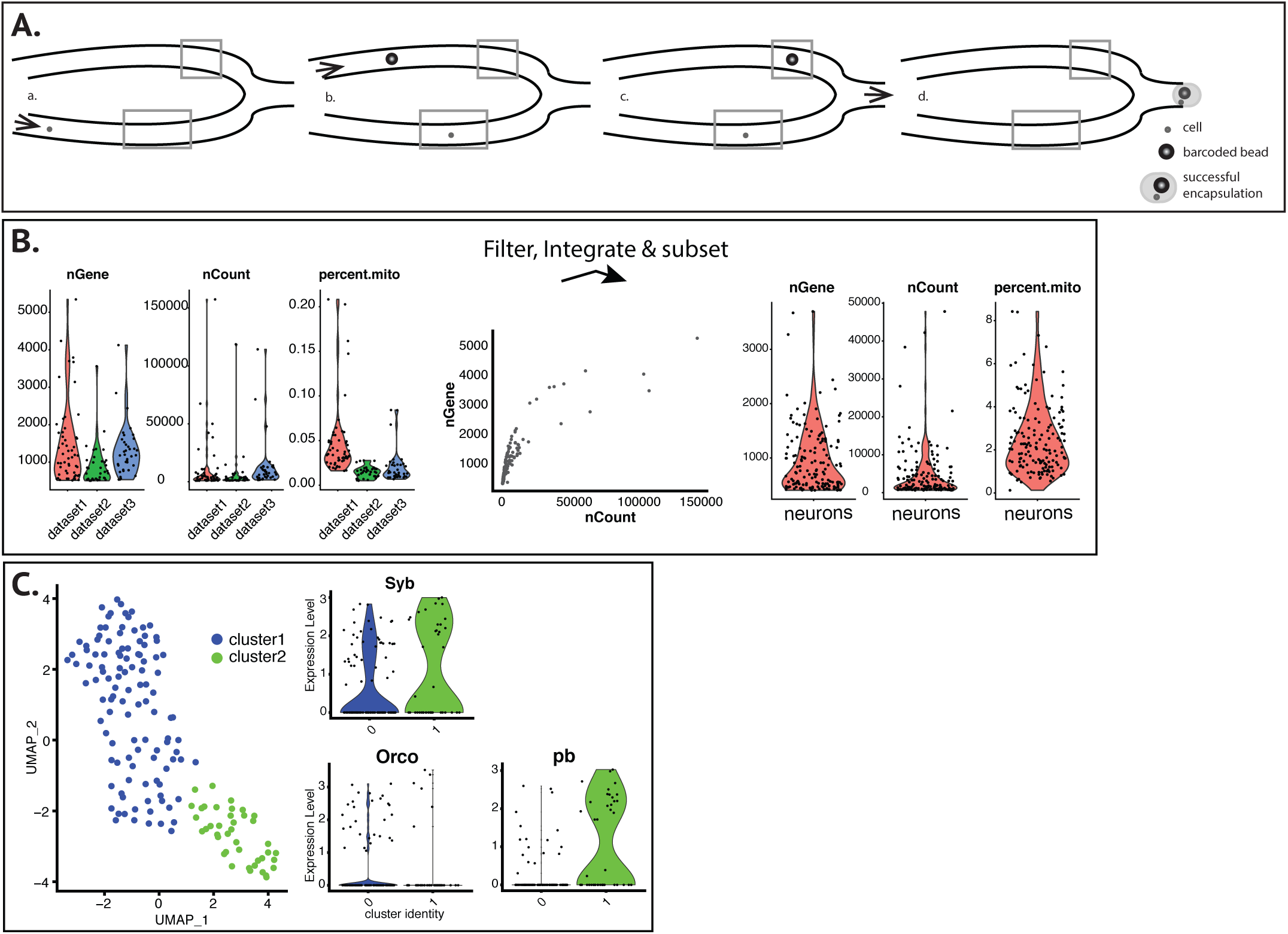
Single-cell RNA sequencing of chemosensory neurons. A. Schematic principle of DisCo (DeterminIStic CO-encapsulation) with optimized co-encapsulation efficiency for minimal cell loss. DropSeq on a microfluidic chip equipped with valves that allow single cells (arrow in a.) to be stalled in a designated region (bottom square, b.) until a corresponding barcoded bead (arrow in b.) reaches a second designated region (upper square, c.) for their concomitant release into an oil droplet (d.). B. Three data sets as biological and technical replicates were merged and filtered to eliminate cells with high gene number (nGene) or high UMIs (nCount) as potential indicator of doublets and cells with high mitochondrial gene expression as indicator of apoptotic cells. Correlation between high gene number and high UMIs (nGene vs nCount) was used for establishing filter thresholds. Integrated data was subset for neurons, resulting in a dataset of 153 cells above 400 genes per cell. C. UMAP embedding in Seurat was colored by cell cluster association of cells. Neuronal marker *Syb* shows expression in both cell clusters, while *Orco* and *pb* show distinct expression roughly corresponding to the identified clusters.

**Supplementary Figure 6:**
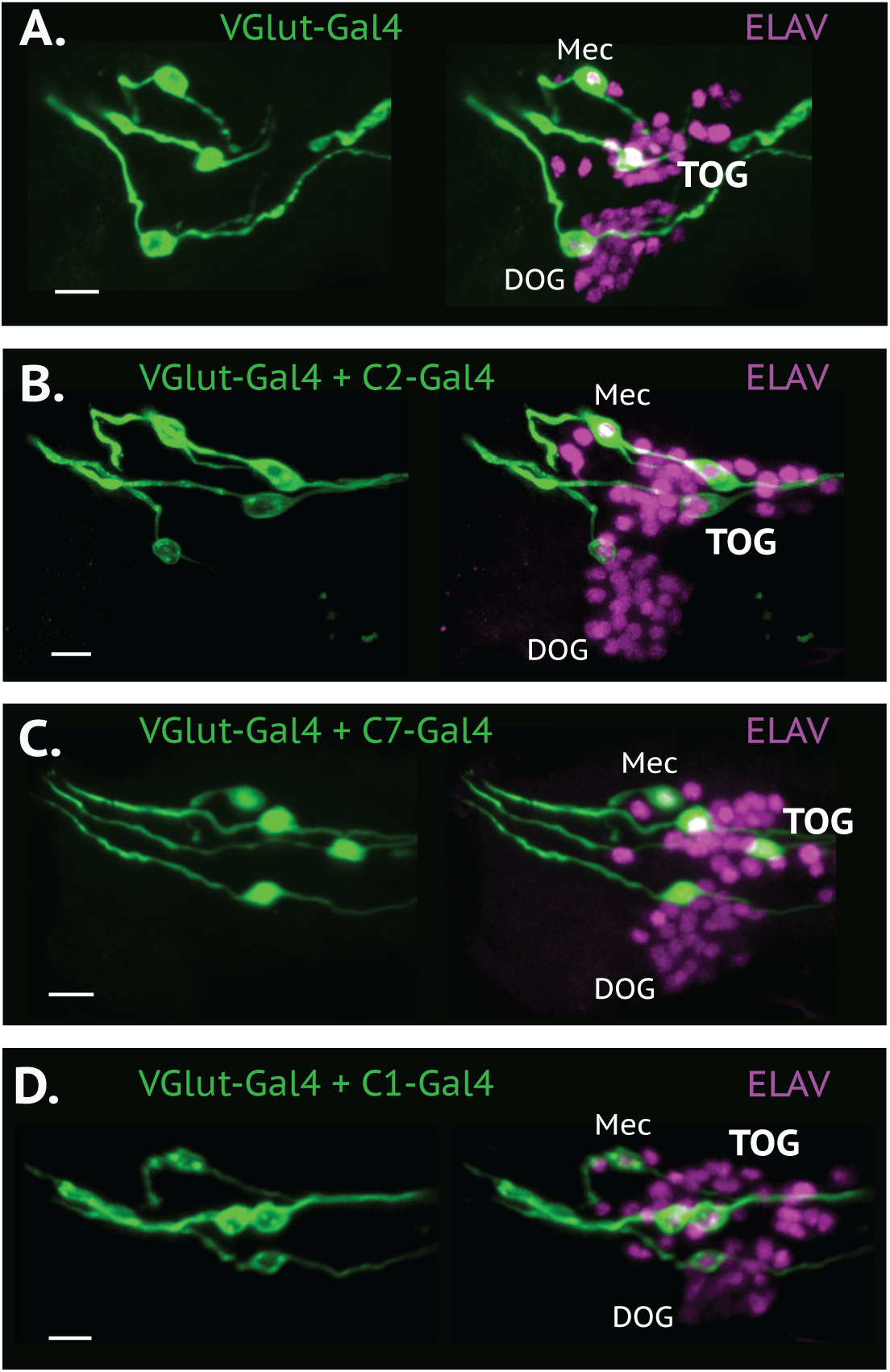
*VGlut*-expressing TOG neuron does not co-localize with C2 or with C7. *VGlut-*positive TOG neuron (**A**.) does not share identity with C2 (**B**.), C7 (**C**.) or C1 (**D**.), using corresponding Gal4 lines and comparing the number of GFP positive cells in separate versus crossed lines. Scale bars, 10 µm.

**Supplementary Figure 8:**
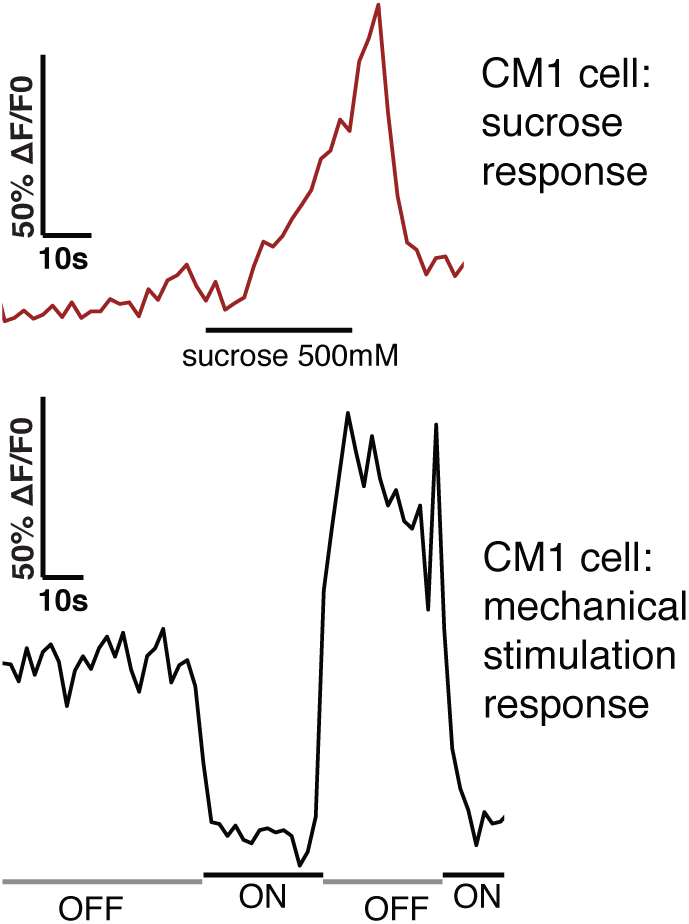
Response to both gustatory and mechanical stimulation in a TOG neuron. CM1 cell identified in a pan-neuronal whole-organ recording responds to sucrose 500mM and is deactivated when the water flow pump is turned on. The cell was already exposed to water inside the microfluidic channel, and therefore the response is likely not determined by water itself but by shear stress on the cell membrane.

**Supplementary Table 1. Chemicals used for taste stimulation.**

**Supplementary File 1. Data referring to Fig.2**.

**Supplementary File 2. Data referring to Fig.4.**

**Supplementary File 3. Data referring to Fig.6C**.

**Supplementary File 4. Whole organ calcium imaging, data processing, Suppl.Fig1 – ImageJ scripts and pipeline instruction files**.

## References

1. Gerber B, Stocker RF. The *Drosophila* larva as a model for studying chemosensation and chemosensory learning: a review. Chemical senses. 2007;32(1):65–89.

2. El-Keredy A, Schleyer M, Konig C, Ekim A, Gerber B. Behavioural analyses of quinine processing in choice, feeding and learning of larval Drosophila. PLoS One. 2012;7(7):e40525.

3. Konig C, Schleyer M, Leibiger J, El-Keredy A, Gerber B. Bitter-sweet processing in larval *Drosophila*. Chem Senses. 2014;39(6):489–505.

4. Kim H, Choi MS, Kang K, Kwon JY. Behavioral Analysis of Bitter Taste Perception in Drosophila Larvae. Chem Senses. 2016;41(1):85–94.

5. Schipanski A, Yarali A, Niewalda T, Gerber B. Behavioral analyses of sugar processing in choice, feeding, and learning in larval Drosophila. Chem Senses. 2008;33(6):563–73.

6. Croset V, Schleyer M, Arguello JR, Gerber B, Benton R. A molecular and neuronal basis for amino acid sensing in the Drosophila larva. Sci Rep. 2016;6:34871.

7. Kudow N, Miura D, Schleyer M, Toshima N, Gerber B, Tanimura T. Preference for and learning of amino acids in larval Drosophila. Biol Open. 2017;6(3):365–9.

8. Stocker RF. The organization of the chemosensory system in *Drosophila melanogaster*: a review. Cell and tissue research. 1994;275(1):3–26.

9. Stocker RF. Design of the larval chemosensory system. Adv Exp Med Biol. 2008;628:69–81.

10. Chandrashekar J, Hoon MA, Ryba NJ, Zuker CS. The receptors and cells for mammalian taste. Nature. 2006;444(7117):288–94.

11. Huang AL, Chen XK, Hoon MA, Chandrashekar J, Guo W, Trankner D, et al. The cells and logic for mammalian sour taste detection. Nature. 2006;442(7105):934–8.

12. Yarmolinsky DA, Zuker CS, Ryba NJ. Common sense about taste: from mammals to insects. Cell. 2009;139(2):234–44.

13. Barretto RPJ, Gillis-Smith S, Chandrashekar J, Yarmolinsky DA, Schnitzer MJ, Ryba NJP, et al. The neural representation of taste quality at the periphery. Nature. 2015;517(7534):373–U511.

14. Smith DV, St John SJ. Neural coding of gustatory information. Curr Opin Neurobiol. 1999;9(4):427–35.

15. Wu A, Dvoryanchikov G, Pereira E, Chaudhari N, Roper SD. Breadth of tuning in taste afferent neurons varies with stimulus strength. Nature Communications. 2015;6.

16. Tomchik SM, Berg S, Kim JW, Chaudhari N, Roper SD. Breadth of tuning and taste coding in mammalian taste buds. Journal of Neuroscience. 2007;27(40):10840–8.

17. Chaudhari N, Roper SD. The cell biology of taste. J Cell Biol. 2010;190(3):285–96.

18. Dahanukar A, Lei YT, Kwon JY, Carlson JR. Two Gr genes underlie sugar reception in Drosophila. Neuron. 2007;56(3):503–16.

19. Fujii S, Yavuz A, Slone J, Jagge C, Song X, Amrein H. *Drosophila* Sugar Receptors in Sweet Taste Perception, Olfaction, and Internal Nutrient Sensing. Curr Biol. 2015.

20. Thorne N, Chromey C, Bray S, Amrein H. Taste perception and coding in Drosophila. Curr Biol. 2004;14(12):1065–79.

21. Liman ER, Zhang YV, Montell C. Peripheral coding of taste. Neuron. 2014;81(5):984–1000.

22. French A, Ali Agha M, Mitra A, Yanagawa A, Sellier MJ, Marion-Poll F. Drosophila Bitter Taste(s). Front Integr Neurosci. 2015;9:58.

23. Zhang YV, Ni J, Montell C. The molecular basis for attractive salt-taste coding in Drosophila. Science. 2013;340(6138):1334–8.

24. Hussain A, Zhang M, Ucpunar HK, Svensson T, Quillery E, Gompel N, et al. Ionotropic Chemosensory Receptors Mediate the Taste and Smell of Polyamines. Plos Biology. 2016;14(5).

25. Lee MJ, Sung HY, Jo H, Kim HW, Choi MS, Kwon JY, et al. Ionotropic Receptor 76b Is Required for Gustatory Aversion to Excessive Na+ in Drosophila. Mol Cells. 2017;40(10):787–95.

26. Tauber JM, Brown EB, Li Y, Yurgel ME, Masek P, Keene AC. A subset of sweet-sensing neurons identified by IR56d are necessary and sufficient for fatty acid taste. PLoS Genet. 2017;13(11):e1007059.

27. Ahn JE, Chen Y, Amrein H. Molecular basis of fatty acid taste in Drosophila. Elife. 2017;6.

28. Ganguly A, Pang L, Duong VK, Lee A, Schoniger H, Varady E, et al. A Molecular and Cellular Context-Dependent Role for Ir76b in Detection of Amino Acid Taste. Cell Rep. 2017;18(3):737–50.

29. Sanchez-Alcaniz JA, Silbering AF, Croset V, Zappia G, Sivasubramaniam AK, Abuin L, et al. An expression atlas of variant ionotropic glutamate receptors identifies a molecular basis of carbonation sensing. Nat Commun. 2018;9(1):4252.

30. Jaeger AH, Stanley M, Weiss ZF, Musso PY, Chan RC, Zhang H, et al. A complex peripheral code for salt taste in Drosophila. Elife. 2018;7.

31. Vosshall LB, Stocker RF. Molecular architecture of smell and taste in *Drosophila*. Annual review of neuroscience. 2007;30:505–33.

32. Scott K, Brady R, Jr., Cravchik A, Morozov P, Rzhetsky A, Zuker C, et al. A chemosensory gene family encoding candidate gustatory and olfactory receptors in *Drosophila*. Cell. 2001;104(5):661–73.

33. Colomb J, Grillenzoni N, Ramaekers A, Stocker RF. Architecture of the primary taste center of *Drosophila* melanogaster larvae. The Journal of comparative neurology. 2007;502(5):834–47.

34. Rimal S, Lee Y. The multidimensional ionotropic receptors of Drosophila melanogaster. Insect Mol Biol. 2018;27(1):1–7.

35. Stewart S, Koh TW, Ghosh AC, Carlson JR. Candidate ionotropic taste receptors in the *Drosophila* larva. Proc Natl Acad Sci U S A. 2015.

36. Alves G, Salle J, Chaudy S, Dupas S, Maniere G. High-NaCl perception in *Drosophila* melanogaster. J Neurosci. 2014;34(33):10884–91.

37. Mast JD, De Moraes CM, Alborn HT, Lavis LD, Stern DL. Evolved differences in larval social behavior mediated by novel pheromones. Elife. 2014;3:e04205.

38. Xu J, Sornborger AT, Lee JK, Shen P. Drosophila TRPA channel modulates sugar-stimulated neural excitation, avoidance and social response. Nat Neurosci. 2008;11(6):676–82.

39. Kwon JY, Dahanukar A, Weiss LA, Carlson JR. Molecular and cellular organization of the taste system in the *Drosophila* larva. The Journal of neuroscience : the official journal of the Society for Neuroscience. 2011;31(43):15300–9.

40. Apostolopoulou AA, Rist A, Thum AS. Taste processing in *Drosophila* larvae. frontiers in integrative Neuroscience. 2015;9.

41. Python F, Stocker RF. Adult-like complexity of the larval antennal lobe of D. melanogaster despite markedly low numbers of odorant receptor neurons. J Comp Neurol. 2002;445(4):374–87.

42. Gendre N, Luer K, Friche S, Grillenzoni N, Ramaekers A, Technau GM, et al. Integration of complex larval chemosensory organs into the adult nervous system of *Drosophila*. Development. 2004;131(1):83–92.

43. Larsson MC, Domingos AI, Jones WD, Chiappe ME, Amrein H, Vosshall LB. Or83b encodes a broadly expressed odorant receptor essential for *Drosophila* olfaction. Neuron. 2004;43(5):703–14.

44. Kreher SA, Kwon JY, Carlson JR. The molecular basis of odor coding in the *Drosophila* larva. Neuron. 2005;46(3):445–56.

45. Fishilevich E, Domingos AI, Asahina K, Naef F, Vosshall LB, Louis M. Chemotaxis behavior mediated by single larval olfactory neurons in *Drosophila*. Current biology : CB. 2005;15(23):2086–96.

46. Mathew D, Martelli C, Kelley-Swift E, Brusalis C, Gershow M, Samuel AD, et al. Functional diversity among sensory receptors in a Drosophila olfactory circuit. Proc Natl Acad Sci U S A. 2013;110(23):E2134–43.

47. Si G, Kanwal JK, Hu Y, Tabone CJ, Baron J, Berck M, et al. Structured Odorant Response Patterns across a Complete Olfactory Receptor Neuron Population. Neuron. 2019;101(5):950–62 e7.

48. van Giesen L, Hernandez-Nunez L, Delasoie-Baranek S, Colombo M, Renaud P, Bruggmann R, et al. Multimodal stimulus coding by a gustatory sensory neuron in Drosophila larvae. Nat Commun. 2016;7:10687.

49. Rist A, Thum AS. A map of sensilla and neurons in the taste system of drosophila larvae. J Comp Neurol. 2017;525(18):3865–89.

50. Bues J, Biočanin M, Pezoldt J, Dainese R, Chrisnandy A, Rezakhani S, et al. Deterministic scRNA-seq of individual intestinal organoids reveals new subtypes and coexisting distinct stem cell pools. BioRxiv. 2020.

51. Su CY, Menuz K, Reisert J, Carlson JR. Non-synaptic inhibition between grouped neurons in an olfactory circuit. Nature. 2012;492(7427):66–71.

52. Zhang Y, Tsang TK, Bushong EA, Chu LA, Chiang AS, Ellisman MH, et al. Asymmetric ephaptic inhibition between compartmentalized olfactory receptor neurons. Nat Commun. 2019;10(1):1560.

53. Python F, Stocker RF. Immunoreactivity against choline acetyltransferase, gamma-aminobutyric acid, histamine, octopamine, and serotonin in the larval chemosensory system of Dosophila melanogaster. J Comp Neurol. 2002;453(2):157–67.

54. Brunet Avalos C, Maier GL, Bruggmann R, Sprecher SG. Single cell transcriptome atlas of the Drosophila larval brain. Elife. 2019;8.

55. Chen TW, Wardill TJ, Sun Y, Pulver SR, Renninger SL, Baohan A, et al. Ultrasensitive fluorescent proteins for imaging neuronal activity. Nature. 2013;499(7458):295–300.

56. Egger B, Boone JQ, Stevens NR, Brand AH, Doe CQ. Regulation of spindle orientation and neural stem cell fate in the Drosophila optic lobe. Neural Dev. 2007;2:1.

57. van Giesen L, Neagu-Maier GL, Kwon JY, Sprecher SG. A microfluidics-based method for measuring neuronal activity in Drosophila chemosensory neurons. Nat Protoc. 2016;11(12):2389–400.

58. Marella S, Fischler W, Kong P, Asgarian S, Rueckert E, Scott K. Imaging taste responses in the fly brain reveals a functional map of taste category and behavior. Neuron. 2006;49(2):285–95.

59. Park J, Carlson JR. Physiological responses of the Drosophila labellum to amino acids. J Neurogenet. 2018;32(1):27–36.

60. Faber DS, Pereda AE. Two Forms of Electrical Transmission Between Neurons. Front Mol Neurosci. 2018;11:427.

61. Biocanin M, Bues J, Dainese R, Amstad E, Deplancke B. Simplified Drop-seq workflow with minimized bead loss using a bead capture and processing microfluidic chip. Lab Chip. 2019;19(9):1610–20.

62. Macosko EZ, Basu A, Satija R, Nemesh J, Shekhar K, Goldman M, et al. Highly Parallel Genome-wide Expression Profiling of Individual Cells Using Nanoliter Droplets. Cell. 2015;161(5):1202–14.

63. Klein AM, Macosko E. InDrops and Drop-seq technologies for single-cell sequencing. Lab Chip. 2017;17(15):2540–1.

64. Satija R, Farrell JA, Gennert D, Schier AF, Regev A. Spatial reconstruction of single-cell gene expression data. Nat Biotechnol. 2015;33(5):495–502.

65. Stuart T, Butler A, Hoffman P, Hafemeister C, Papalexi E, Mauck WM, 3rd, et al. Comprehensive Integration of Single-Cell Data. Cell. 2019;177(7):1888–902 e21.

66. Rusch DB, Kaufman TC. Regulation of proboscipedia in Drosophila by homeotic selector genes. Genetics. 2000;156(1):183–94.

67. Salvaterra PM, Kitamoto T. Drosophila cholinergic neurons and processes visualized with Gal4/UAS-GFP. Brain Res Gene Expr Patterns. 2001;1(1):73–82.

68. Stocker RF. The Organization of the Chemosensory System in Drosophila-Melanogaster - a Review. Cell and Tissue Research. 1994;275(1):3–26.

69. Chu-Wang IW, Axtell RC. Fine structure of the terminal organ of the house fly larva, Musca domestica L. Z Zellforsch Mikrosk Anat. 1972;127(3):287–305.

70. Jeong YT, Oh SM, Shim J, Seo JT, Kwon JY, Moon SJ. Mechanosensory neurons control sweet sensing in Drosophila. Nat Commun. 2016;7:12872.

71. Sanchez-Alcaniz JA, Zappia G, Marion-Poll F, Benton R. A mechanosensory receptor required for food texture detection in Drosophila. Nat Commun. 2017;8:14192.

72. Apostolopoulou AA, Hersperger F, Mazija L, Widmann A, Wust A, Thum AS. Composition of agarose substrate affects behavioral output of *Drosophila* larvae. Frontiers in behavioral neuroscience. 2014;8:11.

73. Kudow N, Kamikouchi A, Tanimura T. Softness sensing and learning in Drosophila larvae. J Exp Biol. 2019;222(Pt 7).

74. Tracey WD, Jr., Wilson RI, Laurent G, Benzer S. painless, a Drosophila gene essential for nociception. Cell. 2003;113(2):261–73.

75. Brown TD. Techniques for mechanical stimulation of cells in vitro: a review. J Biomech. 2000;33(1):3–14.

76. Reiter S, Campillo Rodriguez C, Sun K, Stopfer M. Spatiotemporal Coding of Individual Chemicals by the Gustatory System. J Neurosci. 2015;35(35):12309–21.

77. Hernandez-Nunez L, Belina J, Klein M, Si G, Claus L, Carlson JR, et al. Reverse-correlation analysis of navigation dynamics in larva using optogenetics. Elife. 2015;4.

78. Jisoo Han MC. Comprehensive functional screening of taste sensation in vivo. bioRxiv. 2018.

79. Smith DV, Travers JB. Metric for the Breadth of Tuning of Gustatory Neurons. Chemical Senses & Flavour. 1979;4(3):215–29.

80. Jefferys JG. Nonsynaptic modulation of neuronal activity in the brain: electric currents and extracellular ions. Physiol Rev. 1995;75(4):689–723.

81. Miriyala A, Kessler S, Rind FC, Wright GA. Burst Firing in Bee Gustatory Neurons Prevents Adaptation. Curr Biol. 2018;28(10):1585–94 e3.

82. Saro G, Lia AS, Thapliyal S, Marques F, Busch KE, Glauser DA. Specific Ion Channels Control Sensory Gain, Sensitivity, and Kinetics in a Tonic Thermonociceptor. Cell Rep. 2020;30(2):397–408 e4.

83. Meissner GW, Nern A, Singer RH, Wong AM, Malkesman O, Long X. Mapping Neurotransmitter Identity in the Whole-Mount Drosophila Brain Using Multiplex High-Throughput Fluorescence in Situ Hybridization. Genetics. 2019;211(2):473–82.

84. Xu Y, Qin S, Niu Y, Gong T, Zhang Z, Fu Y. Effect of fluid shear stress on the internalization of kidney-targeted delivery systems in renal tubular epithelial cells. Acta Pharm Sin B. 2020;10(4):680–92.

85. White CR, Frangos JA. The shear stress of it all: the cell membrane and mechanochemical transduction. Philos Trans R Soc Lond B Biol Sci. 2007;362(1484):1459–67.

86. Al-Anzi B, Tracey WD, Jr., Benzer S. Response of Drosophila to wasabi is mediated by painless, the fly homolog of mammalian TRPA1/ANKTM1. Curr Biol. 2006;16(10):1034–40.

87. Singh RN, Singh K. Fine-Structure of the Sensory Organs of *Drosophila-Melanogaster* Meigen Larva (Diptera, Drosophilidae). International Journal of Insect Morphology & Embryology. 1984;13(4):255–73.

88. Schindelin J, Arganda-Carreras I, Frise E, Kaynig V, Longair M, Pietzsch T, et al. Fiji: an open-source platform for biological-image analysis. Nat Methods. 2012;9(7):676–82.

89. Parslow A, Cardona A, Bryson-Richardson RJ. Sample drift correction following 4D confocal time-lapse imaging. J Vis Exp. 2014(86).

90. Team RC. R: A language and environment for statistical computing. R Foundation for Statistical Computing, Vienna, Austria. URL https://www.R-project.org/. 2017.

91. Wickham H. FR, Henry L. and Müller K. dplyr: A Grammar of Data Manipulation. R package version 0.7.6. https://CRAN.R-project.org/package=dplyr. 2018.

92. Team R. RStudio: Integrated Development for R. 2015.

93. Wickham H. Reshaping Data with the reshape Package. Journal of Statistical Software. 2007;21(12):1–20.

94. Wickham H. ggplot2: Elegant Graphics for Data Analysis. 2009.

95. Wickham H. SD, RStudio. scales: Scale Functions for Visualization. 2019.

96. Picelli S, Bjorklund AK, Reinius B, Sagasser S, Winberg G, Sandberg R. Tn5 transposase and tagmentation procedures for massively scaled sequencing projects. Genome Res. 2014;24(12):2033–40.

97. Macosko E.Z. GM, McCarroll S.A. Drop-Seq Laboratory Protocol version 3.1. 2015.

98. Dobin A, Davis CA, Schlesinger F, Drenkow J, Zaleski C, Jha S, et al. STAR: ultrafast universal RNA-seq aligner. Bioinformatics. 2013;29(1):15–21.

