## Supplementary material for "Multimodal and multisensory coding in the *Drosophila* larval peripheral gustatory center": SupplTable 1.pdf

**Supplementary Table 1: Chemicals used for taste stimulation.**

| Tastant or taste group | Compound | Associated taste (humans) | Concentration | Group final Concentration |
|---|---|---|---|---|
| **Individual tastant series 1** | Sucrose | Sweet | 500mM | - |
| | Denatonium benzoate | Bitter | 10mM | - |
| | Valine | Amino acid/Bitter | 100mM | - |
| | NaCl | Salty | 1M | - |
| | Citric Acid | Sour | 100mM | - |
| **Individual tastant series 2** | Sucrose | Sweet | 500mM | - |
| | Quinine | Bitter | 5mM | - |
| | Arginine | Amino acid/Bitter | 100mM | - |
| | NaCl | Salty | 100mM | - |
| | Citric Acid | Sour | 100mM | - |
| **Group sugars Monosaccharides** | Fructose | Sweet | 100mM | 500mM |
| | Glucose | Sweet | 100mM | |
| | Arabinose | Sweet | 100mM | |
| | Mannose | Sweet | 100mM | |
| | Galactose | Sweet | 100mM | |
| **Group sugars Disaccharides** | Sucrose | Sweet | 100mM | 500mM |
| | Trehalose | Sweet | 100mM | |
| | Maltose | Sweet | 100mM | |
| | Lactose | Sweet | 100mM | |
| | Cellobiose | Sweet | 100mM | |
| **Group bitter DSoTC** | Denatonium benz. | Bitter | 1mM | |
| | Sucrose octaacetate | Bitter | 1mM | |

| | | | | |
|---|---|---|---|---|
| | Theophylline | Bitter | 10mM | 22mM |
| | Coumarin | Bitter | 10mM | |
| **Group bitter QLSC** | Quinine | Bitter | 1mM | 13mM |
| | Lobeline | Bitter | 1mM | |
| | Strychnine | Bitter | 1mM | |
| | Caffeine | Bitter | 10mM | |
| **Group Amino acids A** | Valine | Bitter | 10mM | 60mM |
| | Leucine | Bitter | 10mM | |
| | Isoleucine | Bitter | 10mM | |
| | Methionine | Bitter | 10mM | |
| | Tryptophan | Bitter | 10mM | |
| | Cysteine | Bitter | 10mM | |
| **Group Amino acids B** | Alanine | Sweet | 10mM | 41mM |
| | Phenylalanine | Bitter | 10mM | |
| | Glycine | Sweet | 10mM | |
| | Proline | Sweet | 10mM | |
| | Tyrosine | Bitter | 1mM | |
| **Group Amino acids C** | Arginine | Bitter | 10mM | 50mM |
| | Lysine | Salty-Bitter | 10mM | |
| | Aspartic acid | Umami | 10mM | |
| | Glutamic acid | Umami | 10mM | |
| | Histidine | Bitter | 10mM | |
| **Group Amino acids D** | Serine | Sweet | 10mM | 40mM |
| | Threonine | Sweet | 10mM | |
| | Asparagine | Neutral | 10mM | |

| | | | | |
|---|---|---|---|---|
| | Glutamine | Sweet | 10mM | |
| **Group high salt** | NaCl | Salty | 500mM | 1M |
| | KCl | Salty | 500mM | |
| **Group low salt** | NaCl | Salty | 25mM | 50mM |
| | KCl | Salty | 25mM | |
